# Haplotype independence contributes to evolvability in the long-term absence of sex in a mite

**DOI:** 10.1101/2023.09.07.556471

**Authors:** Hüsna Öztoprak, Shan Gao, Nadège Guiglielmoni, Alexander Brandt, Yichen Zheng, Christian Becker, Kerstin Becker, Viktoria Bednarski, Lea Borgschulte, Katharina Atsuko Burak, Anne-Marie Dion-Côté, Vladislav Leonov, Linda Opherden, Satoshi Shimano, Jens Bast

**Affiliations:** Institute of Zoology, University of Cologne; Cologne, Germany; Department of Ecology and Evolution, University of Lausanne; Lausanne, Switzerland; Cologne Center for Genomics (CCG), Medical Faculty, University of Cologne; Cologne, Germany; Genomics & Transcriptomics Laboratory, Biologisch-Medizinisches Forschungszentrum, Heinrich-Heine-Universität Düsseldorf; Düsseldorf, Germany; Département de Biologie, Université de Moncton; Moncton, Canada; A.N. Severtsov Institute of Ecology and Evolution, Russian Academy of Sciences; Moscow, Russia; Science Research Center, Hosei University; Tokyo, Japan

## Abstract

Some unique asexual species persist over time and contradict the consensus that sex is a prerequisite for long-term evolutionary survival. How they escape the dead-end fate remains enigmatic. Here, we generated a haplotype-resolved genome assembly based on a single individual and collected genomic data from worldwide populations of the parthenogenetic diploid oribatid mite *Platynothrus peltifer* to identify signatures of persistence without sex. We found that haplotypes diverge independently since the transition to asexuality at least 20 my ago. Multiple lines of evidence indicate disparate evolutionary trajectories between haplotypic blocks. Our findings imply that such haplotypic independence can lead to non-canonical routes of evolvability, helping some species to adapt, diversify and persist for millions of years in the absence of sex.

**One-Sentence Summary:** Functionally different chromosome sets in an asexual mite species showcase a survival strategy spanning millions of years.

## Main Text

Some oribatid mite species are rare evolutionary anomalies, as they maintain effective purifying selection and persist and diversify over millions of years in the absence of sex (*1*–*4*). Oribatid mites are diverse (>10,000 species), small (150-1400 μm), mainly soil-living decomposers that were among the first arthropods to colonize land during the Devonian (*5, 6*). Notably, a high number of species in this animal order (10 %) reproduce via parthenogenesis, of which many radiated and form diverse phylogenetic clades (*7, 8*). Such evolutionary exceptions are invaluable because understanding the peculiarities for success without sex will help to identify the adaptive value and constraints of sex vice-versa (*9*). To date, however, the population-genomic signatures of persistence without sex, i.e. how peculiar genome dynamics of ancient asexuals might contribute to maintaining effective selection, adaptation and diversification remain largely unknown (*1, 3, 10*).

### A phased genome assembly to identify signatures of ancient asexuality

One of the most iconic old asexual species is the diploid oribatid mite *Platynothrus peltifer* (C.L. Koch, 1893; **Fig. 1A**). Previous work suggested a transition to asexuality tens of millions of years ago, likely predating the separation of Europe and North America (*2*). The cytological mechanism for asexuality is a form of automictic thelytoky during which recombination is restricted to chromosome ends, maintaining heterozygosity and resulting in ‘effective clonality’ (*11*–*13*). *Platynothrus peltifer* has a generation time of one year and produces only female offspring in lab rearings, and extremely rare spanandric males are infertile (*14, 15*).

**Fig. 1.**
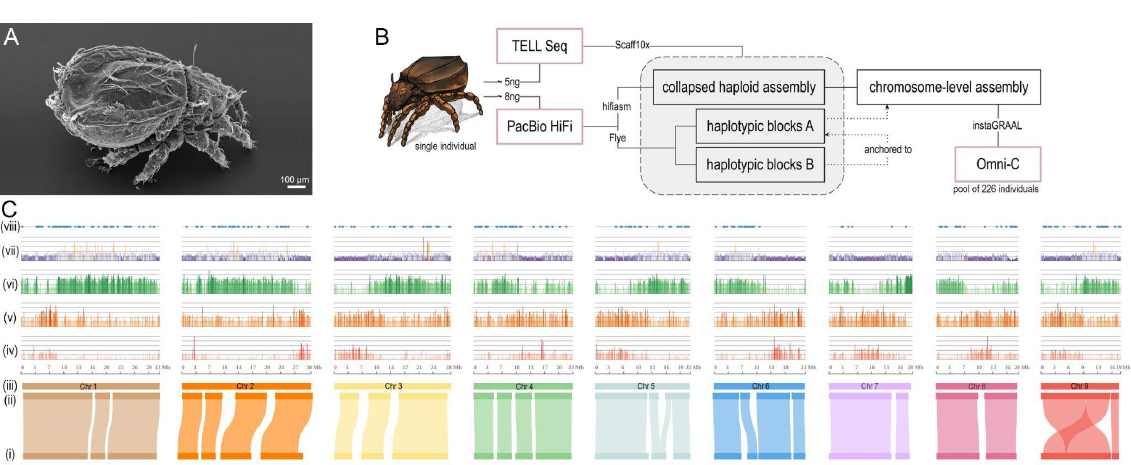
Assembly strategy and characteristics of the *Platynothrus peltifer* genome. (**A**) *P. peltifer* electron microscopy picture. Scale bar indicates 100 μm. (**B**) Schematic workflow of single-individual sequencing and assembly. (**C**) Genomic properties of the aligned haplotypic blocks (i) A and (ii) B, anchored to the (iii) nine pseudochromosomes, for which densities of 100 kb blocks of (iv) coding sequences, (v) gene number, (vi) transposable elements, (vii) heterozygosity percentage, and (viii) horizontally transferred gene locations are shown along the genome.

To reveal the genomic substrate for persistence without sex, we first generated a chromosome-scale reference genome from a natural population and additionally resolved haplotypes of *P. peltifer* from a single individual, sampled in Germany (**Fig. 1**). In short, PacBio HiFi long reads and TELL-Seq linked reads stemming from one diploid individual were assembled into haplotype-collapsed scaffolds that were ordered to chromosome-scale with chromosome conformation capture (Hi-C) data from pooled individuals of the same population. The same long-read and linked-read data were used to generate a haplotype-resolved assembly; following, phased haplotypic blocks were anchored to each other and the chromosome-scale assembly (**Fig. 1B, fig. S1)**. The chromosome-level genome assembly is of high completeness and contiguity and spans 219 Mb comprising nine chromosomes with 24,932 annotated genes (**Fig. 1C, fig. S2, fig. S3, table S1**). The haplotypes comprise 63 blocks B anchored to 44 blocks A, representing 92.7 % (203 Mb) of the genome being phased (**Fig. 1C, table S2**). The genome size estimate via flow cytometry closely matches the assembly size (6 % difference: 232 Mb vs 219 Mb) and together with *k*-mer spectra ploidy analyses suggest no ancient whole genome duplication (**fig. S4, supplementary text**). Additionally, we assembled the complete mitochondrial genome from the same individual (**fig. S5**).

### Haplotype dynamics and divergence across worldwide populations

After the transition from sexuality to asexuality, spontaneous mutations are predicted to occur independently on each haplotype in a diploid, ‘effectively clonal’ asexual. Over time, the absence of sex, recombination and gene flow should thus manifest into increasing intra-individual heterozygosity, which drives the divergence and independent evolution of haplotypes. Consequently, if the transition to asexuality occurred a considerable amount of time ago, such that different populations separated after, haplotypes should be more diverged within individuals than between individuals from different populations, generating haplotype trees mirroring each other’s topology. Such independent evolution of haplotypes is the strongest evidence for obligate asexual reproduction over long time periods (aka, the ‘Meselson effect’) (*16*). However, despite this straightforward prediction, empirical evidence remains equivocal and the only strong evidence for independent haplotype evolution in animals stems from an oribatid mite species (*1*).

To investigate haplotypic independence associated with ancient asexuality in *P. peltifer*, we sequenced five individuals per population from German (Dahlem), Italian (Montan), Russian (Moscow Oblast), Japanese (Yamanashi) and Canadian (Moncton) populations (**Fig. 2A**). Genetic divergence clusters individuals by their geographical locations (**Fig. 2B**). Next, to identify haplotypic differences among individuals and populations, we analyzed genetic diversity patterns over the whole nine chromosomes. Overall dynamics of mean heterozygosity between haplotypes within individuals are similar among populations, with shared regions of high and low heterozygosity (**Fig. 2C**). This suggests differences in purifying selection or mutation rate heterogeneity among the various chromosomal regions, shared by all populations. German, Italian and Russian (European) populations feature similar levels of mean individual heterozygosity of 1.6 % to 1.8 % along chromosomes. As expected under asexuality, individuals within these populations also share a large proportion of heterozygous sites, with Italian individuals sharing most (**fig. S6**). The mean heterozygosity of individuals from Canadian (1.4 %) and Japanese (1.0 %) populations is consistently lower compared to the other populations, but shows strikingly similar heterozygosity dynamics along chromosomes (**Fig. 2C**). These differences are thus not the consequence of large distinct stretches of loss of heterozygosity (LOH) that can occur under some non-clonal forms of asexuality (*17*). Individuals within these populations moreover share a lower proportion of heterozygous sites, especially Canadian individuals (**fig. S6**). These findings suggest considerable differences in processes that can affect the overall divergence of haplotypes among populations, such as the rate of spontaneous mutations increasing heterozygosity, the rate of gene conversion removing heterozygosity and/or suggest independent and more recent transitions to asexuality for the Japanese and Canadian populations.

**Fig. 2.**
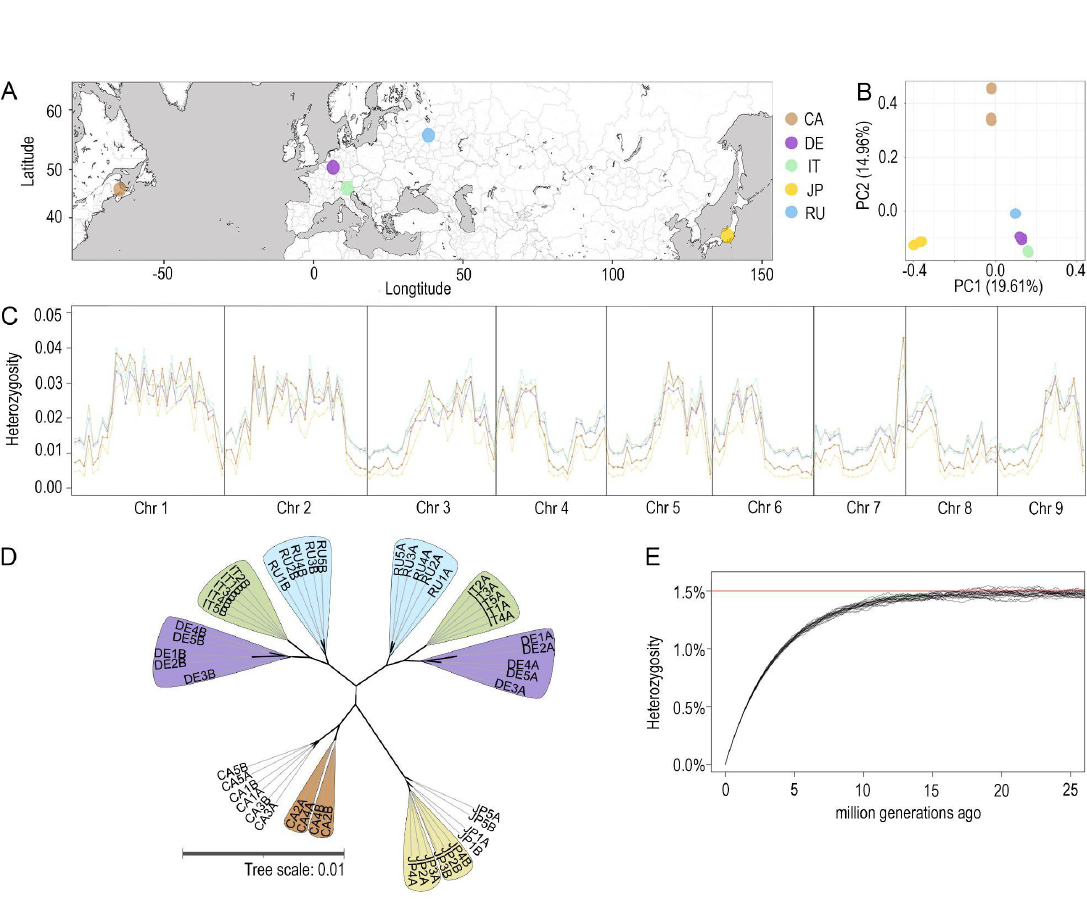
Haplotype dynamics of worldwide *Platynothrus peltifer* populations suggest independent haplotype evolution for at least 20 my under asexuality of the DE-IT-RU lineages. (**A**) Sampling spots of the *P. peltifer* populations worldwide. Background gray shading indicates water bodies. Abbreviations: CA: Canada, DE: Germany, IT: Italy, JP: Japan, RU: Russia, color code kept throughout the figure. (**B**) Principal component analysis (PCA) of genomic population data visualizing 34.6 % of the total variability. (**C**) Mean heterozygosity between haplotypes within individuals for each population along the nine chromosomes, separated into 1 Mb genomic blocks. (**D**) Unrooted consensus haplotype tree showing the Meselson effect, i.e., complete separation of haplotypes A and B with mirror topologies, among *P. peltifer* populations from Germany, Russia and Italy. Bold branches indicate consensus supports > 96 %. (**E**) Simulations of increased divergence between haplotypes over time under gene conversion track length of 1000 bp, reaching a plateau after 20 million generations, using biological parameters of *P. peltifer*.

### Haplotypes evolve independently under asexuality since 20 million years ago

We generated haplotype-specific trees using phased data to identify if the transition to asexuality occurred considerable amounts of time before the separation into different populations (**Fig. 2D, fig. S7, table S3**). As expected under ancient asexuality, a perfect split of the two haplotypes displaying mirror topologies of populations could be identified, including all individuals of European populations, representing the ‘Meselson effect’ (**Fig. 2D**). Contrastingly, haplotype trees of Japanese and Canadian populations lack such clear separation. While some individuals display a shared separation of haplotypes as expected under asexuality, this is confined to each respective population (**Fig. 2D**). Taken together, heterozygosity patterns are corroborated by haplotype topologies (**Fig. 2C, D, fig. S7**) and suggest an ancient transition to asexuality for the ancestor lineage of the European populations and a more recent independent loss of sex for both the Japanese and Canadian populations. Heterozygosity dynamics along chromosomes are very similar in all populations and imply conserved synteny in the ancestral lineage for all transitions (**Fig. 2C**). Thus, while likely not sharing the same transition event, the ancestor of all *P. peltifer* populations was a closely related lineage, indicating comparable ages of asexuality or very conserved genome evolution. Although the mechanism of transition to asexuality is unknown, hybridization is unlikely as it can not generate the shared haplotype differences among the European populations. Moreover, hybridization would entail substantial genomic changes and thus can not explain the similar heterozygosity patterns for all populations (*18*). As remnants of *Wolbachia* endosymbiont can be detected in the *P. peltifer* genome, the transition to asexuality might have happened via reproductive manipulation of this endosymbiont (**fig. S8**). This type of transition often occurs via gamete duplication and results in fully homozygous lineages (*25*). Taken together, and given that haplotypic divergence of the Canadian and Japanese populations are considerably lower, transitions to asexuality in *P. peltifer* likely occurred via such a mechanism that substantially removes heterozygosity.

Following, we estimated the age of asexuality (**Fig. 2E**). Haplotypic divergence (i.e., heterozygosity) under asexuality over a substantial amount of time can be used to infer the age of asexuality. Heterozygosity gain over time involves the combined effects of accumulating differences between haplotypes via novel mutations (μ), and decreases via gene conversion events. Consequently, we first estimated the spontaneous mutation rate of *P. peltifer* by sequencing mothers and their eggs from the German population and measured μ = 2.05×10^−9^ (**supplementary text**). Second, we estimated gene conversion to occur with a minimum track length of 500 bp in the German *P. peltifer* (**fig. S9**). Simulations of these combined effects show that contrary to common assumptions, heterozygosity reaches an equilibrium value that is independent of gene conversion track lengths over time (**Fig. 2E, fig. S10**). Using these biological parameters of *P. peltifer*, about 20×10^6^ generations are necessary to attain the mean 1.5 % heterozygosity equilibrium for *P. peltifer* (**Fig. 2E**). Given a generation time of one generation per year for *P. peltifer*, and assuming that the pre-asexuality heterozygosity for *P. peltifer*’s ancestor was substantially lower and that heterozygosity currently is at equilibrium, the European *P. peltifer* lineage has reproduced asexually since at least 20 million years. Moreover, these findings again suggest that the Japanese and Canadian populations are younger as they have not yet reached equilibrium.

### Adaptation and evolutionary innovation in the long-term absence of sex

Asexual organisms are typically regarded as evolutionary dead-ends because linkage of loci should result in decreased efficiency of purifying selection and reduced adaptive potential (*19*). Contrary to evidence for these predictions in some younger asexual arthropods (*20, 21*), asexual species of oribatid mites can maintain effective purifying selection (*3*). However, the genomic substrate for their unique ability to evolve and adapt in the long-term absence of sex remains unknown. Hence, we identified different evolutionary dynamics associated with the exceptional independent evolution of haplotypes that could contribute to adaptation and evolutionary novelty. This is exemplified in the haplotype-resolved genome of German *P. peltifer* (**Fig. 3**) encompassing allelic diversification and functionalization, the impact of horizontally transferred genes, and transposable element activity.

**Fig. 3.**
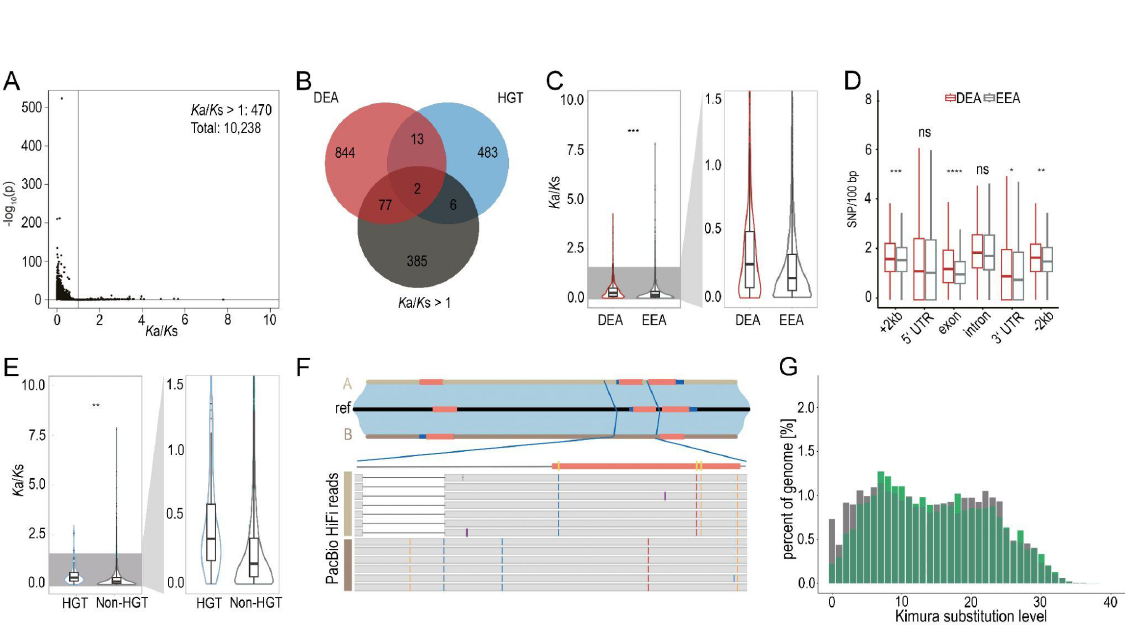
Independent haplotypes contribute to evolvability. (**A**) Pairwise comparison of the *K*_a_*/K*_s_ distribution for allelic genes. (**B**) Venn diagram showing numbers of differentially expressed alleles (DEA), horizontally transferred genes (HGT), and genes with *K*_a_*/K*_s_ > 1. (**C**) Violin plots show levels of *K*_a_*/K*_s_ of differentially expressed alleles and equivalently expressed alleles (EEA), highlighting *K*_a_*/K*_s_ < 1.5. (**D**) Boxplots calculated for SNPs/100 bp for genic and neighboring regions, including the 5’ and 3’ untranslated regions (UTR), exons, introns, and up-/ and downstream regions (+2 kb and -2 kb respectively) for differentially (DEA) and equivalently (EEA) expressed alleles. (**E**) Violin plots show levels of *K*_a_*/K*_s_ of HGTs and non-HGTs, highlighting *K*_a_*/K*_s_ < 1.5. (**F**) Differentially expressed HGT (*Ppr_hap0_g20454*) as an example of the molecular changes contributing to divergent haplotypes. The two haplotypic blocks A and B differ in the number of exons (red bars). The highlighted region shows HiFi reads supporting allelic divergence (brown bars) by a deletion (white bar) and three non-synonymous substitutions (yellow stripe) in the second exon (red bar). Colored stripes in the reads indicate non-reference base substitutions. (**G**) Transposable element divergence landscape of the largest homologous haplotypic blocks of chromosome five (indicated by gray and green). Statistics were calculated using the Student’s t-test between two groups (DEAs (n=879) and EEAs (n=9,359) and between HGTs (n=62) and non-HGTs (n=10,176)) (*: p-values < 0.05, **: < 0.01, ***: < 0.001).

The decoupling and independent evolution of haplotypes after the transition to asexuality in *P. peltifer* (**Fig. 2**) mimics duplications of the whole genome with genes representing quasi-homeologs. Gene duplications are a major source of evolutionary innovation and allow access to novel phenotypes, as they can provide a novel function while preserving the old function of one duplicate and systematically increase mutational robustness (*22, 23*). To test this prediction, we first analyzed differences in genetic variation between haplotypes that show a mean heterozygosity of 2.4 % (**Fig. 1C**). Among 11,029 allelic 1:1 orthologs, the vast majority differ from each other by non-synonymous (88.0 %) and/or synonymous (96.8 %) variants. Approximately 10 % gained or lost a stop codon and about 11 % showed frameshift, and nearly 15 % non-frameshift insertions and/or deletions (**fig. S11**). Non-synonymous to synonymous divergence (*K*_a_*/K*_s_) analysis suggests allele-specific direction of selection with 4.6 % (470/10,238) of alleles showing signs of strong positive selection (*K*_a_*/K*_s_ > 1), potentially indicating rapid evolution (**Fig. 3A**). Over time, diverging alleles (similar to duplicated genes) might acquire novel functionalities and expression patterns. Hence, we also tested whether differentially expressed alleles (DEAs) contribute to functional diversification. About 9.1 % (936) of alleles are differentially expressed and show elevated *K*_a_*/K*_s_ values compared to equivalently expressed alleles (EEAs) (**Fig. 3B, C**). Specifically, upstream and exonic regions of DEAs showed elevated variant densities compared to EEAs, unlike introns and 5’ UTRs, suggesting functional and adaptive diversifications of alleles as well as associated transcription factors (**Fig. 3D**). Up- or down-regulation of DEAs is not haplotype-specific and can be different for each allele. These DEAs are enriched for basic cell functions, such as ribosomes, translation, and protein production (**fig. S12**).

Another mechanism providing novel traits to organisms is horizontal transfer of pre-existing genes (*24*). Horizontal gene transfers (HGTs) represent 2.0 % (504) of *P. peltifer* genes, which is within the typical range of asexual animals (**Fig. 1C, Fig. 3B**) (*25*). These HGT stem mainly from bacteria, but also fungi, plants and metazoans. Of these HGTs, 92.9 % (468) are expressed and 61.5 % (319) contain intronic regions, indicating adjustments to functional integration in the host genome. From the 62 allelic 1:1 HGT copies that show signs of diversification, 25.2 % are differentially expressed, again suggesting selection of diverging HGT alleles (**Fig. 3B, E, F**). These HGTs might have arrived before the transition to asexuality, but divergent haplotypes likely contributed to novel (sub-) functions, similarly to overall alleles. This is why we next identified HGTs that were incorporated after the transition to asexuality and before potential gene conversion events, i.e. HGTs that reside only on one haplotype (**fig. S13**). Of the 33 ‘orphan HGTs’, 19 (57.6 %) are expressed and 16 (48.5 %) contain at least an intron, which is less compared to allelic HGTs, indeed indicating a more recent arrival with less time to adjust to the host background. Notably, HGT functional annotations, including differentially expressed and orphan HGTs, suggest the involvement in processes to digest plant cell walls (e.g. Glycosyl hydrolases). Further, HGT genes of the UGT family may contribute to pesticide resistance, indicating a contribution to the mite’s ecology as soil-living decomposers (**fig. S14, S15, S16, supplementary data**). Overall, the molecular underpinnings driving diverging haplotypes under asexuality, specifically allelic differentiation of DEAs and HGTs, involve in addition to single nucleotide variants, (non-) frameshift insertions and deletions and stop gains/losses, most pronounced in HGT alleles (**Fig. 3F, fig. S11**).

Another possible agent of evolvability are transposable elements (TEs). They proliferate throughout genomes independently of the host cell cycle and their activity is often deleterious (*26*). While TEs likely evolve to be more benign in asexual genomes compared to sexuals (*4, 26*), TE activity can occasionally be beneficial by accelerating evolution and by rewiring regulatory networks (*27, 28*). We identified TEs and their haplotype-specific activity in the *P. peltifer* reference genome. Transposable elements comprise 27 % of the genome assembly and show signs of recent and past activity as emphasized by Kimura substitution levels (**fig. S17, table S4**). Moreover, TE density distributions suggest effective selection against TE insertion within genes, in concordance with previous results (**fig. S18**) (*4*). We detect noticeable differences in historical TE activity between the largest phased haplotypic blocks of chromosomes (**Fig. 3G, fig. S19**). Interestingly, pronounced differences occur from 6 %-12 % TE copy divergence, suggesting an increase in TE activity for one haplotype and a decrease in the other over 29 to 59 my ago. This correlates with the transition to asexuality of the *P. peltifer* lineage over 20 my ago. Very recent activity is largely restricted to one haplotype and, similar to haplotypic alleles, suggests divergence of one haplotype and more conservation of the other in TE activity and content.

Overall, our analyses suggest conservation of one allelic copy (or haplotype) and relaxed selection and/or functional adaptive divergence in the other, similar to innovation via gene duplication (*22*). This suggests that evolution of genes and regulatory regions via haplotypic divergence, modification of pre-existing genes (HGTs) and haplotype-specific activity of transposable elements can lead to increased evolvability via evolutionary innovation and robustness.

### Summary

In summary, we provide support for at least 20 million years of asexual evolution in multiple geographically separated populations of the oribatid mite *P. peltifer*. Several lines of evidence indicate that haplotypic independence provides the substrate for evolvability and evolutionary innovation. Thus far, the best evidence for the Meselson effect and ancient asexuality in parthenogenetic animals only exists in oribatid mites (this study and also (*1*)), suggesting that the rarity of long-term asexuals might be explained by the requirement for haplotypic independence as a source for novelty to adapt and diversify in the absence of sex.

## Acknowledgments

Mark Maraun and Katja Wehner for electron microscopy picture of *Platynothrus peltifer*, Marcel Solbach for the *P. peltifer* drawing. Mélanie Jean for collection of mites in Canada. Nobuhiro Kaneko (Fukushima University) provided information about his previous study site as a habitat for *P. peltifer*.

This work was supported by the DFG Research Infrastructure West German Genome Center (407493903) as part of the Next Generation Sequencing Competence Network (project 423957469). Computational support of the Zentrum für Informations- und Medientechnologie, especially the HPC team (High Performance Computing) at the Heinrich-Heine University is acknowledged.

## Funding

German Research Foundation Emmy Noether grant BA 5800/4-1 (JB). NSERC discovery grant RGPIN-2019-05744 (AMDC)

## Author contributions

Conceptualization: JB

Resources: HÖ, VL, AMDC, SS

Methodology: JB, HÖ, SG, NG, AB, YZ, KAB, VB

Investigation: HÖ, LO, CB, KB

Visualization: HÖ, SG, NG, AB, YZ, VB, LB

Funding acquisition: JB

Project administration: HÖ, JB

Supervision: JB, HÖ, NG, SG

Data curation: NG, SG

Formal Analyses: HÖ, SG, AB, NG, YZ, VB, LB, KAB

Writing – original draft: JB, HÖ, with input from all authors

Writing – review & editing: JB, HÖ

## Competing interests

Authors declare that they have no competing interests.

## Data and materials availability

All code for the analyses can be found at https://github.com/TheBastLab/Ppr_evolution. The genome, together with its annotation is available at NCBI and the reads at the SRA under Bioproject PRJNA1003031 (haplotypic blocks A: PRJNA1019947; haplotypic blocks B: PRJNA1019950)

## Notes

### Competing Interest Statement

The authors have declared no competing interest.

### Summary of Updates

Author affiliations updated. Spelling errors fixed. Description of Figures updated to include the full genus name of Platynothrus peltifer. An additional sentence added to clarify that the age estimation is a minimum estimate.

