## Supplementary Materials and Text for "Haplotype independence contributes to evolvability in the long-term absence of sex in a mite"

**The PDF file includes:**

Materials and Methods

Supplementary Text

Figs. S1 to S19

Tables S1 to S4

References

**Other Supplementary Materials for this manuscript include the following:**

Data S1

### Materials and Methods

Sample preparation and sequencing

**Sample collection.** All mite individuals were sampled from natural populations in Germany, Italy, Canada, Japan and Russia. German samples were collected in a coniferous forest in Dahlem (50.39010 N, 6.57162 E) in a 30 cm² square. Italian samples were collected at Montan Southern Tyrol (46.33514 N, 11.29728 E) in a 30 cm² square. Japanese soil and leaf litter was sampled near a larch forest in Yamanashi (35.5449 N 138.2403 E) and sent to Germany for extraction. Canadian samples were collected from 46.03570 N, -64.79896 E and 46.14482 N, -64.76913 E. Specimens of interest were sent to Germany. Russian samples were isolated from a spruce forest in Moscow Oblast (56.02522 N, 38.43074 E). Specimens were isolated out of leaf litter by heat gradient extraction (*29*). *Platynothrus peltifer* was identified morphologically using (*30*) and molecularly confirmed by cytochrome oxidase I (CO1) sequencing.

**Sample preparation.** Specimens were starved for more than a week and cleansed with a brush in distilled water, distilled water with detergent (fit GmbH, Zittau, Germany), and incubated in NaClO 0.05% (DonKlorix; CP GABA GmbH, Hamburg, Germany) and ethanol 70% for 30 seconds each and rinsed in distilled water again.

**High-molecular-weight *g*DNA extraction (for ultra-low input).** For high-molecular-weight (HMW) single-individual DNA extraction, we established a modified salting out protocol (*31*). In short: A single individual was submerged in TNES buffer and flash-frozen in liquid nitrogen. The sample was then homogenized using a sterile pestle. After adding Proteinase K, the sample was incubated for at least 1h. Next, yeast tRNA was added, followed by NaCl and 96% ethanol. DNA purification was conducted and the sample was left to homogenize overnight. DNA concentration was measured using Qubit Fluorometer v. 4 with the Qubit dsDNA HS Assay kit (Thermo Fisher Scientific, Waltham, MA).

**Cytochrome oxidase I sequencing**. The cytochrome oxidase subunit 1 (CO1; ~700 bp) region was amplified using the universal primers LC01490F/HC02198R (Folmer et al.,

1994). The PCR reaction was run with 1 ng of DNA, 2x Thermo Scientific DreamTaq Green PCR Master Mix, 1 µM forward and 1 µM reverse primer, and ddH20 to fill until 25 µl. Amplification was conducted under the following conditions: denaturation at 95°C for 5 min, 35 cycles at 95°C for 30 s, 45°C for 30 s and 72°C for 1 min, and final extension at 72°C for 6 min. PCR products were purified by adding 1 U/ml of Exonuclease and 0.3 U/ml FastAP to 8 µl PCR product then heated for 30 min at 37°C, and subsequently for 20 min at 85°C. For sequencing, the Big dye Terminator Cycle sequencing Kit and an ABI PRISM automatic sequencer were used at the Cologne Center for Genomics (CCG; Cologne, Germany). To molecularly verify the samples, sequences of Cytochrome oxidase I with >99% similarity to *P. peltifer* were identified as such.

**DNA quality.** HMW *g*DNA sample was assessed at the Genomics & Transcriptomics Laboratory (GTL; Düsseldorf, Germany) using the Agilent Femto Pulse system. HMW *g*DNA with an OD260/280 ratio of approximately 1.8 to 2.0 and fragment size above 15 kb was selected for sequencing.

**Long-read and linked-read sequencing for reference assembly**. Single-individual HMW DNA was sequenced using two technologies: PacBio HiFi (SMRTbell® Libraries from Ultra-Low DNA Input) and TELL-seq (TELL-seq™ WGS library)TM both with ultra-low DNA input of at least 5 ng. Library preparation and sequencing of the SMRTbellTM templates were conducted by the GTL. PacBio HiFi sequencing yielded 34.7 Gb of reads with an N50 of 15 kb. In addition, TELL-seq libraries were constructed and sequenced by CCG using a TELL-Seq WGS Library Prep Kit (Universal Sequencing Technology, Carlsbad, CA) and yielded 69 Gb of reads.

**Omni-C sequencing.** To construct the library for Omni-C adult individuals were sampled from two spots in the coniferous forest in Dahlem, Germany in close vicinity. Previous analyses on cytochrome oxidase I from these two spots suggest two distinct mitochondrial lineages to be present in both of these spots. 226 whole adult specimens were flash-frozen with liquid $N_2$, crushed, vortexed, and the Omni-C Proximity Ligation Assay protocol for insects & marine invertebrates was followed. The library was sequenced on Illumina NovaSeq 6000 by Novogene (Cambridge, United Kingdom), which generated $92.7x10^6$ pairs of 150-bp reads.

**Linked-read sequencing for populations.** In addition to the reference individual, four individuals of the same population were sequenced using Illumina TELL-seq linked paired-end reads only. For details see supplementary data.

**RNA sequencing.** Total RNA was extracted from ten adult individuals using TRIzol reagent treated with DNase I within a Direct-zolTM RNA MicroPrep kit (Zymo Research, Irvine, CA). An RNA-seq library was constructed by the CCG using TruSeq Stranded Total RNA with Ribo-Zero Globin and $43\text{x}10^6$ pairs of 100-bp reads were sequenced.

***k*-mer analyses.** 27-mers in the HiFi reads were analyzed using KAT v2.4.2 (*32*) with the modules kat hist and kat gcp (default parameters). Ploidy was further investigated using kmc v3.2.1 with parameters -k 27 -ci 1 -cs 10000 and Smudgeplot (v0.2.5) (*33*) with default parameters.

Chromosome-level collapsed and phased assemblies

***De novo* genome assembly.** HiFi reads were assembled using hifiasm (v0.16.1-r375) (*34*) with default parameters into initial collapsed haploid contigs, and using Flye (v2.9) (*35*) with default parameters into phased contigs.

**TELL-seq scaffolding.** Barcoded TELL-seq linked Illumina reads were generated and corrected from BCL raw data using tell-read (v1.0.2). The barcodes were formatted to 10X Genomics standard by using ust10x (v1.0.2) from TELL-seq conversion_tool. The collapsed haploid contigs (Hap0: 211 Mb, N=33) were scaffolded using Scaff10X (v4.2) (github.com/wtsi-hpag/Scaff10X) with formatted TELL-seq reads and using arguments -longread 1 -gap 100 -matrix 2000 -reads 10 -score 10 -edge 50000 -link 8 -block 50000. Haplotype-resolved contigs were scaffolded using Scaff10X with parameters -longread 1 -gap 100 -matrix 2000 -reads 10 -score 10 -edge 50000 -link 8 -block 50000.

**Omni-C scaffolding.** Omni-C reads were cleaned using TrimGalore (v0.6.5) (github.com/FelixKrueger/TrimGalore) with parameters '-j 30 -q 30 --fastqc --paired' and mapped to the draft haploid assembly using hicstuff (v3.1.1) and bwa (v0.7.15) (*36*) with parameters --enzyme 100 --iterative --aligner bwa. instaGRAAL (v0.1.6 no-opengl branch) (*37*) was run with parameters --level 5 --cycles 100. The scaffolds were curated with instaGRAAL-polish to reduce misassemblies and add 10 Ns in gaps. 99.75% of the assembly were anchored to nine chromosome-level scaffolds which were selected as chromosome candidates for subsequent analyses.

**Gap filling and polishing.** HiFi reads were mapped to the haploid assembly using minimap2 with parameters '--secondary=no --MD -ax asm20' and then sorted and indexed using Samtools (v1.11) with default parameters. Gaps in the collapsed scaffolds Hap0 were filled using TGS-GapCloser (v1.1.1) (*38*) with the parameters '-tgstype pb --minmap_arg -x asm20 --ne' and the final haploid (Hap0) assembly was polished using HyPo (v1.0.3) (*39*) with the PacBio HiFi reads.

**Haplotype scaffolding.** Primary and alternative haplotypes were separated using minimap2 and purge_dups (v1.2.5) (*40*), and are subsequently designated as haplotypic blocks A and B. Haplotypes were reciprocally scaffolded using RagTag (v2.1.0) (*41*) with the other haplotype as reference.

**Anchoring of haplotypes.** Haplotypic blocks A were mapped against the collapsed assembly (Hap0) using minimap2 (v2.24-r1122) with parameter -x asm5. Haplotypic blocks B were mapped against haplotypic blocks A. Haplotype-resolved scaffolds were attributed to chromosome candidates of Hap0 for which they had the higher number of residue matches.

Assembly evaluation

**Completeness.** For the chromosome-level assembly (Hap0), ortholog completeness was assessed using the tool Benchmarking Universal Single-Copy Orthologs (BUSCO v.5.0.0) (*42*) against the Arthropoda odb10 lineage (1,066 orthologs) and the Arachnida odb10 lineage (2,934 orthologs). *k*-mer completeness was evaluated using KAT (v2.4.2) and the module kat comp with default parameters.

**Omni-C contact map.** Omni-C reads were mapped to Hap0 using bwa and hicstuff as previously described. The contact map was generated using the module hicstuff view with the parameter -b 500.

**Contaminants.** PacBio HiFi reads were mapped to the final Hap0 scaffolds using minimap2 (v2.24-r1122) with parameters -ax map-hifi and the mapped reads were sorted with SAMtools (v1.11) (*43*, *44*). Hap0 was aligned against the nucleotide database using the Basic Local Alignment Search Tool (BLAST v2.6.0) (*45*) with parameters -outfmt "6 qseqid staxids bitscore std sscinames scomnames" -max_hsps 1 -evalue 1e-25. The outputs of minimap2, BLAST, and BUSCO (against the Arachnida odb10 lineage) were provided as input to Blobtools2 (*43*).

Genome annotation

**Repeat and transposable element annotation and masking**. A repeat library including transposable elements (TEs) was built using EDTA (v1.9.6) (*46*) with parameters '--sensitive 1 --anno 1'. TEs were annotated and classified to the superfamily level using the FasTE pipeline (*47*). The hardmasked assemblies were converted to softmasked assemblies using bedtools (v2.26.0) with mask mode.

**RNA-seq mapping.** Adapter sequences were removed from RNA-seq raw reads using TrimGalore (v0.6.5) with parameters '-j 30 -q 30 --fastqc --paired'. Trimmed reads were mapped to the softmasked reference haploid assembly Hap0 and the two haplotypic blocks assemblies haplotypic blocks A and haplotypic blocks B using STAR (v2.5.1a) (*48*) with parameters '--readFilesCommand zcat --outSAMtype BAM SortedByCoordinate --outSAMstrandField intronMotif --outFilterIntronMotifs RemoveNoncanonical'.

**Gene Structure Prediction**. Trinity (v2.1.1) was employed to assemble the RNA-Seq reads *de novo*, and the resulting 61,067 transcripts were aligned to the assemblies using PASA (v2.5.2). Mapped reads were provided to StringTie (v2.2.0) to predict transcripts. Then TransDecoder (https://github.com/TransDecoder/TransDecoder/wiki; Last accessed November 14, 2018) was used to find the Open Reading Frames (ORF) for each gene. BRAKER2 (v2.1.6) was used to predict gene structures with RNA-seq evidence. These results were provided to EVidenceModeler (v1.1.1) with weight parameters 'ABINITIO_PREDICTION AUGUSTUS 4; TRANSCRIPT assembler-database.sqlite 7; OTHER_PREDICTION transdecoder 8' to select genes. Finally, PASA was used to refine UTR regions. The final annotation was converted into protein sequences using gffread (v0.12.1).

**Functional annotation.** We utilized Eggnog-mapper (v2.1.6) with protein sequences and generated KEGG (Kyoto Encyclopedia of Genes and Genomes) pathway annotations. Furthermore, we employed InterProScan (version 5.61-93.0) with protein sequences to generate Reactome pathway annotations. The Gene Ontology (GO) annotation was created by combining the results of Eggnog-mapper (v2.1.6) and InterProScan (version 5.61-93.0).

<u>Genome dynamics</u>

**TE abundance and landscapes.** Quality-cleaned genome-wide repeat abundances were generated by FasTE. To generate the TE divergence landscape a custom R script was created to rename the TE superfamilies after Wicker-classification (*49*) and adjust the data to plot the TE divergence Landscape with the 'Plot Kimura Distance' R script for the Hap0 and the largest haplotypic blocks per chromosome. For more information on the size of regions see table S2.

**TE Density.** The TE Density pipeline (*50*) was applied to generate the TE Density plots, which are defined as TE-occupied base pairs in a given window, on the collapsed assembly of *P. peltifer*.

**Gene synteny.** Protein sequences of Hap0 and the haplotypic blocks were aligned using BLAST v2.6.0 with parameters -evalue 1e-10 -outfmt 6. In order to find blocks of gene synteny, MCScanX (commit 97e74f4) was run between Hap0, haplotypic blocks A and haplotypic blocks B with default parameters. Collinearity was visualized with SynVisio (*51*) with a minimum match score >= 3950.895. Heterozygosity between the haplotypes was estimated using Mash (v2.3) (*52*).

<u>Mitochondrial genome assembly and annotation</u>

TELL-seq reads were trimmed using TrimGalore and assembled using MitoFinder v1.4.1 with the mitochondrial genome of *Steganacarus magnus* as seed (*53*, *54*). The assembly was annotated using MiTFi v0.1 (*55*) and then using MITOS (*56*), MITOS2 (*57*)(both available at http://mitos2.bioinf.uni-leipzig.de/index.py), ARWEN v1.2.3 (*58*) and tRNAscan-SE v2.0 (*59*) to find genes and tRNAs that could not be found by MiTFi. tRNAscan-SE was run with COVE cutoff set to -20. Geneious (Biomatters Ltd.) was used to manually curate annotations and find consensus predictions. tRNAs were selected based on their predicted secondary structure and their minimum free energy, computed using the RNAfold web server (http://rna.tbi.univie.ac.at/) (*60*).

Spontaneous mutation rate estimation

**Sample preparation.** For spontaneous *de novo* mutation rate estimation without rearings and mutation accumulation lines, best practice was followed by sequencing parents and offspring (as in (*61*, *62*)). *Platynothrus peltifer* individuals were sampled from the same population as the reference genome individual (Dahlem, Germany) in early June 2021. For three individual mothers (M1, M2, and M3) their 'offspring daughters', i.e. eggs, were extracted after removing the genital plates. Eggs were cleansed in NaClO 0.05% (DonKlorix; CP GABA GmbH, Hamburg, Germany) with a brush to remove tissue from their mothers. For M1 one daughter (D1), for M2 one daughter (D2), and for M3 three daughters (D3, D4, and D5) could successfully be analyzed.

**Sequencing.** To avoid PCR-induced read errors and to get highly accurate reads, the 'NEBNext Ultra II FS DNA Library Prep Kit for Illumina' (New England Biolabs, Ipswich, USA) in combination with UMI Adaptor 'NEBNext Multiplex Oligos for Illumina (Unique Dual Index UMI Adaptors DNA Set 1)' (New England Biolabs, Ipswich, USA) was chosen. These libraries were multiplexed and whole-genome sequenced on an Illumina NovaSeq 600 generating paired-end 150 bp reads. This yielded a total of $811.6 \times 10^6$ reads with a mean of $102.1 \times 10^6$ for the mothers (min $100.3 \times 10^6$, max $103.0 \times 10^6$) and a mean of $100.9 \times 10^6$ (min $81.2 \times 10^6$, max $111.0 \times 10^6$) for the eggs.

**Identifying *de novo* mutations (DNM).** Reads were demultiplexed using UMI-tools extract (v.1.1.2). Adapters were trimmed using TrimGalore (v0.6.5) with parameters -q 30 -a AGATCGGAAGAGCACACGTCTGAACTCCAGTCA -a2 AGATCGGAAGAGCGTCGTGTAGGGAAAGAGTGT --fastqc --paired. Reads were mapped against the hardmasked reference genome of *P. peltifer* using the Burrows-Wheeler Alignment tool *bwa mem* (v0.7.17-r1198-dirty) and filtered with the Genome Analysis ToolKit (GATK v4.1.9). Duplications were removed with UMI-tools (v 1.1.2). Potential variant sites were called and filtered using GATK (v4.1.9) following an in-house pipeline (all scripts and parameters are outlined in detail at the github). In short, variants were filtered by read depth, phred-scaled likelihood of the genotype, allele depth and homozygous to heterozygous mutations. Additionally, following common practice (*62*, *63*) only candidate sites that were homozygous in mothers and heterozygous in daughters were considered as possible *de novo* mutations. Identified mutation site candidates were manually checked in Integrative Genomics Viewer (IGV v2.8.13) under the following conditions: mothers were filtered for a major allele frequency of 0.95 and the daughters for a minor allele frequency of 0.2. Alternatively, a major allele frequency of 0.9 for the mothers and a minor allele frequency of 0.3 for the daughters also passed the filter. Candidate sites found in all sibling daughters were removed as false positives resulting from erroneous genotype assignment in the mother (see Supplementary Data Table 3 ).

**Spontaneous mutation rate.** To calculate the mutation rate per generation, the number of sites for which a new variant could have been detected, i.e. the 'callable genome' was identified. The callable genome includes all homozygous sites in the 3 mothers, similarly filtered as the DNMs. All positions with the variant sites are included in the previously called GVCF-file by GATKs HaplotypeCaller. Each mother's GVCF-file was filtered in read depth (DP), genotype (GT), phred-scaled likelihood of the genotype (PL), and allele depth (AD): DP was filtered like above, sites with DP between 28 to 124 were accepted; for GT, all homozygous sites from the mother were selected; PL for sites must have fulfilled $PL_{second\ most\ likely} - PL_{first\ most\ likely} < 120$; only sites where AD supported one allele were kept. For details see github.

Population genetic analyses

**Variant identification**. Single nucleotide polymorphisms (SNPs) were used to investigate population dynamics within populations of asexual *P. peltifer* from Germany, Russia, Italy, Canada and Japan. Phased population data were generated by mapping trimmed raw-reads generated by TELL-read (v1.0.3) using *bwa mem* (v0.7.15) with default parameters to the collapsed soft masked haploid reference genome (Hap0). The resulting alignment was sorted using samtools (v1.11) and duplications were removed using Picard MarkDuplicates (v2.26.2 Broad Institute). Coverage was calculated with samtools flagstat. Following, variants were called using the Genome Analysis ToolKit (GATK v.4.1.9.0) pipeline. GVCFs were produced using HaplotypeCaller and then merged using CombineGVCFs. Variants were detected with

GenotypeGVCFs. SNPs were selected with SelectVariants. PLINK (v 1.9) and VCFtools (0.1.15) were used to perform PCA analysis.

**Population data coverage filter**. To compare population statistic metrics among the different populations (DE, RU, IT, JP, CA), it is required that the respective reference genome sites are matched, i.e. aligned. Some genomic regions may have only one haplotype being successfully sequenced or mapped, which may lead to artifactual underestimation of heterozygosity and false signals of homozygosity. This becomes apparent with regions mapping with half coverage of the overall median mapped coverage. To mitigate this issue, for each individual the site-wise coverage was filtered, so that only sites that have 75% or higher of the genomic median coverage (for that individual) are used for the population statistics. There is high variation of coverage distribution among individuals (median ranging from 28 in IT3 to 124 in IT1, though most are between 70 and 100) which makes this individual-based coverage filtering necessary.

**Basic population statistics.** The German reference genome is divided into contiguous 1 Mb blocks for each chromosome, with a total of 220 blocks. Within each block, we calculated the heterozygosity for each individual (see github for scripts used). Because calculation of $\theta\pi$ or $\theta w$ requires using only sites that passed the coverage filter for every individual and thus the amount of data is too small and potentially biased, we used an alternative method to quantify deviation from sexual equilibrium: the 'unshared-to-shared ratio' by comparing two individuals from the same population at a time, rx = (H_u1+H_u2)/(2*H_sh) where H_u1 is the number of sites that are heterozygous only in the first individual, H_u2 heterozygous only in the second individual, and H_sh heterozygous in both. For each population, all ten possible pairs are used and the mean value is used. This ratio can show if the heterozygosity and site frequency distribution is more similar to a sexual one (closer to 2) or a clonal one (closer to 0), and is comparable in interpretation to the heterozygosity-to-$\theta\pi$ ratio (which is 1 for a sexual equilibrium population and 2 for a long-term clonal one).

**Empirical gene conversion estimation.** To compare empirical data with the simulated gene conversion data (see simulations), we tallied the distribution of 'homozygosity stretches', i.e., the distance between two consecutive heterozygous sites, from each individual in the German sample. To minimize the effect of mapping artifacts, only stretches of 1000bp or shorter are counted, and the stretch is required to have its mean coverage above the filter threshold. The distribution is calculated separately for each individual and is compared with simulated data under different gene conversion track lengths.

Haplotype-specific analyses

**Parallel divergence of haplotypes.** Quality trimmed paired-end resequencing reads were mapped against haplotypic block assemblies haplotypic blocks A and haplotypic blocks B simultaneously (competitive mapping) and split according to which haplotype they mapped best to using bbsplit (bbmap v38.63; (*64*)). Reads with ~> 5 noncontiguous substitutions were discarded (minratio=0.9). Reads that mapped to both haplotypic block assemblies were kept (ambiguous2=split) and merged with each the set of split reads mapping best to haplotypic blocks A and to the set of split reads mapping best to haplotypic blocks B per individual. This was done to avoid biasing the analysis towards regions that are phaseable, and hence highly heterozygous, in most populations and individuals.

The split sets of resequencing reads (haplotypic blocks A+ambiguous reads and haplotypic blocks B+ambiguous reads) were mapped to the collapsed, softmasked genome assembly Hap0 using bwa v0.7.17 and sorted using samtools v 1.15.1 (as described above). Optical and sequencing duplicates were removed using picard MarkDuplicates v2.26.2 (Broad Institute). Variants were called for each split set of resequencing reads, i.e. each haplotype, separately using gatk v4.2.6.1 HaplotypeCaller, CombineGVCFs and GenotypeGVCFs as described above. Indels, multiallelic sites, sites with a quality $< 20$, genotypes with a depth below 10 and heterozygous genotypes were removed using bcftools. Heterozygous genotypes could result from sequencing error, paralogs and incomplete phasing: either due to the absence of a region from one of the two haplotypic block assemblies such that reads from both haplotypes map to the haplotypic block assembly that is present, or due to low divergence of haplotypes yielding ambiguous mapping results.

Variants were applied to the collapsed assembly using bcftools consensus thereby generating one consensus genome per haplotype per individual. Missing sites and genotypes were applied as Ns leaving a 50 sequence whole haplome alignment (2 haplotypes * 5 individuals * 5 populations). To determine regions that were in-phase, i.e. no phase switching possible within a region, scaffolds of haplotypic blocks assembly haplotypic blocks A were aligned to the collapsed assembly using D-Genies and standard parameters (see Table S3) (*65*). Per chromosome the longest alignment block was extracted from the haplome alignment using geneious (*66*) and subdivided into 1 Mio Mb long bins using msa_split (*67*). Best fitting ML trees were reconstructed for each bin using IQ-tree (version 1.6.1) (*68*) with implemented model testing and 1000 bootstrap replicates. A majority rule consensus tree (i.e. comprises clades that are present in at least 50% of trees) was constructed using geneious (see Figure 2; Figure S9).

**Horizontal gene transfer (HGT).** To detect horizontally transferred genes from different organisms, a previously developed pipeline (https://github.com/reubwn/hgt) (*69*) was utilized, which applies several lines of evidence for detecting HGT candidates (HGTc). In short, DIAMOND (v0.9.21.122) BLASTP was used in 'sensitive' mode to align the protein sequences from the reference and two haplotypic block assemblies to the UniRef90 database. Then taxids information from the NCBI Taxonomy database and three measures were employed to identify putative candidate HGTs.

The three measures including the (a) HGT Index (hU) using the best-hit bitscores from the BLASTP results and defined '(best-hit bitscore for OUTGROUP) - (best-hit bitscore for INGROUP)'; (b) Alien Index (AI) based on E-values also from the BLASTP results to evaluate HGT defined by 'log10 ((best-hit E-value for INGROUP) + 1e-200) - log10((best-hit E-value for OUTGROUP) + 1e-200)'(*70*) and then (c) Consensus Hit Support (CHS) using the sum of bitscores across all hits, and the HGTc must be designated as outgroup and the designation of rest hits >=90%. IQ-tree (version 1.6.1) was used to plot gene phylogenies.

**Orphan HGTs.** To detect HGTc that are only found on one of the two haplotypic blocks ('orphan HGTs'), the protein sequence of the HGTs from one haplotypic block was searched in the other haplotypic block respectively. The potential orphan HGTs were additionally analyzed for coverage, as true missing HGTs should exhibit substantially lower coverage as only one copy (instead of two) and single copy genes from BUSCO results are sequenced. Mapped reads count was divided by gene length for normalization.

Additionally, the orphan HGT candidates were checked for syntenic neighboring gene content.

**Variation calling and annotation.**

Using PacBio HiFi reads from the reference individual, we mapped them to the Hap0 genome using minimap2 (version 2.24-r1122). Subsequently, the resulting BAM file was sorted using

Samtools (version 1.11). The identical pipeline as previously employed was then applied and the SNP annotated by ANNOVAR (version 2020-06-07 23:56:37 -0400).

**Orthology and Ka/Ks.** The protein sequences of annotated genes from the haplotypic blocks and collapsed assemblies were used for detecting the single-copy genes between haplotypes. Among these three, pairwise comparisons were made by OrthoFinder (v 2.5.4) (*71*) to detect single-copy orthogroups genes with default parameters on the haplotypic blocks and collapsed assemblies. The 1:1 orthologous genes between haplotypic blocks A and haplotypic blocks B were additionally filtered by the 1:1 orthology results from the haplotypic blocks A to Hap0 and haplotypic blocks B to Hap0. Then the ParaAT (v 2.0) (*72*) integrated KaKs_Calculator (v 1.2) was employed to calculate the Ka/Ks value for the haplotypic orthologous single-copy genes pairs.

**Differential expression of alleles (DEA)**. The cleaned RNA-Seq reads used for annotation were mapped to the two softmasked haplotypic block assemblies haplotypic blocks A and haplotypic blocks B using STAR (v2.5.1a) with parameters '--readFilesCommand zcat --outSAMtype BAM SortedByCoordinate --outSAMstrandField intronMotif --outFilterIntronMotifs RemoveNoncanonical --outFilterMismatchNmax 3 --outFilterMismatchNoverLmax 0.1 --outFilterMismatchNoverReadLmax 0.5 '.
Differential expression of alleles (DEA) between the two haplotypic blocks haplotypic blocks A and haplotypic blocks B were performed with GFOLD V1.1.4 (*73*). The GFOLD value ≥ | 0.2 | and the RPKM>1 of all samples were set as cutoff values (The GFOLD value could be considered as a reliable log2 fold change). All identified DEAs were further submitted for functional enrichment with R package clusterProfiler (v4.6.2), using all RPKM>1 single-copy genes as the background with the 'enricher' function.

<u>Wolbachia</u>

Previously published *Wolbachia* genomes on NCBI were utilized ASM1658442v1 as a query. To identify candidate genes integrated into the mite genome, we employed the DIAMOND (v. 0.9.21) Blastp tool. Genes exhibiting an identity of over 40% were deemed potential candidates for integration into the mite genome. The synteny was plotted by using NGenomeSyn (v1.41) (*74*).

Simulations

**Single-lineage-based long-term simulation.** Considering a lack of exchange of genetic material between individuals in a cloning species, long-termed evolution can be efficiently simulated by tracing the sequences of a pair of haplotypes (i.e., an individual) over time. To determine the long-term effects of cloning and gene conversion on heterozygosity, we simulated a genomic block as a sequence of 5 million sites. Each site can be in a state of homozygosity (represented by value 0) or heterozygosity (represented by value 1).

At the beginning (generation 0), all 5 million sites are set to 0. Mutations occur with a rate of $2\times\mu$ events per site per generation. When it occurs, the site is set to 1 regardless of the original state (in other words, back-mutations from heterozygotes to homozygotes are considered negligible). Gene conversion occurs with a rate of $2\times r_{GC}$ events per site per generation. When it occurs, $L_{GC}$ consecutive sites (starting from a random site) are set to 0, also regardless of the original state. The "2×" in both mutations and gene conversions represents the fact that both types of events can initiate in one or the other haplotype.

The mutation rate is set to $\mu = 2.05\times10^{-9}$ (as empirically estimated, see above). The rate of gene conversion is set up in such a way that $r_{GC}\times E(L_{GC}) = \mu/0.015$, which would lead to an equilibrium heterozygosity of 1.5% (to match the average empirical divergence between haplotype of the populations). We set the mean gene conversion track length, $E(L_{GC})$ as a variable with values 50, 100, 200, 500, 1000, 2000 and 5000; and the $L_{GC}$ for each gene conversion event is either constant or geometrically distributed. For each of the 7×2 parameter combinations, 20 replicates are run. Each replicate runs for $5\times10^{7}$ generations. To speed up the computation, every 100 generations are completed in the same loop, with the mutations occurring first and gene conversion second. Every 5000 generations, the number of heterozygous sites in the 5 Mb sequence is recorded. From $2\sim5\times10^{7}$ generations, we also record the distribution of homozygous stretch lengths (numbers of consecutive homozygotes) every $1\times10^{5}$ generations.

Genome size estimation

**Flow Cytometry.** To estimate the genome size of *P. peltifer*, a flow cytometry was utilized following the method of (*75*), supported by personal communication with J. Spencer Johnston. *Drosophila melanogaster* (1 C = 175 Mb) was used as the reference. In short, animals were frozen at -80°C and cut on dry ice. The head of *D. melanogaster* was removed and ground with 15 strokes (1 per second) of a B pestle in 10 µL of Galbraith buffer in a 1 mL Dounce homogenizer, after which 990 µL buffer were added and homogenized. For the mite, the anterior portion of the proterosoma was cut off (as mites do not have distinct 'heads'), typically containing blood cells, muscle and neural cells that will yield 2C nuclei. This part of the mite was similarly ground in 10µL Galbraith buffer, after which 440 µL budder was added. Following, 50 µL of the *D. melanogaster* mix was added to the mite cells and homogenized. The mix containing sample and reference was filtered through a 45-µm nylon mesh and 20 µL of Propidium iodide (50 µg/mL) was added homogenized. The sample was then incubated at 4°C for 2 h prior to running with the Accuri C6 Flow Cytometer (BD Biosciences, Erembodegem, Belgium). To compare fluorescence signals, the sample and reference were run both separately and together to check for 2C and 4C peak signals. The flow cytometry was run slowly and stopped after a minimum of 1000 counts in the 2C peaks. Software gates were used for cleaning and filtering the data. The data was deemed good quality if the peaks were separated and the 4C had the double mean FL3-A value as the 2C peak. Several replicates were run and the best quality peak chosen as result.

### Supplementary Text

#### Genome size estimation with flow cytometry

The relative genome size of *Platynothrus peltifer* was estimated to be 232 Mb. For *P. peltifer* the mean FL3-A for the 2C peak was 395 and for the 4C peak 789. For *D. melanogaster* 298 and 596 respectively. The genome size estimate of flow cytometry matches somewhat closely the cleaned, chromosome-scale genome assembly size (6% difference: 232 Mb vs 219 Mb). Note that flow cytometric estimates for mites are typically associated with some minor uncertainties, as there is no clear head given and only 'few' cells can be extracted and stained. Additionally, *D. melanogaster* genome sizes are variable to some degree among individuals and strains. Following, the genome assembly is a very suitable representation of the full *P. peltifer* genome.

#### Mutation rate estimations

In total, two *de novo* spontaneous mutations were identified in daughters compared to mothers based on the filtered and carefully curated variants. These two mutations were found at chromosome 2 in position 5257052 and at chromosome 4 in position 19592709 in D2.

To determine the overall mutation rate, the number of *de novo* mutations is divided by twice (as the species is diploid) the sum of all callable sites from replicate daughters taken together.

$$\mu = \frac{Number\ of\ de\ novo\ mutations}{(2\times\ callable\ genome)} \rightarrow \mu = \frac{2}{(2\times\ 488{,}839{,}965)} = 2.05 * 10 - 9$$

The estimated spontaneous mutation rate is thus $2.05x10^{-9}$ per generation for *P. peltifer* and is thus within the range of *Daphnia* and *Drosophila*.

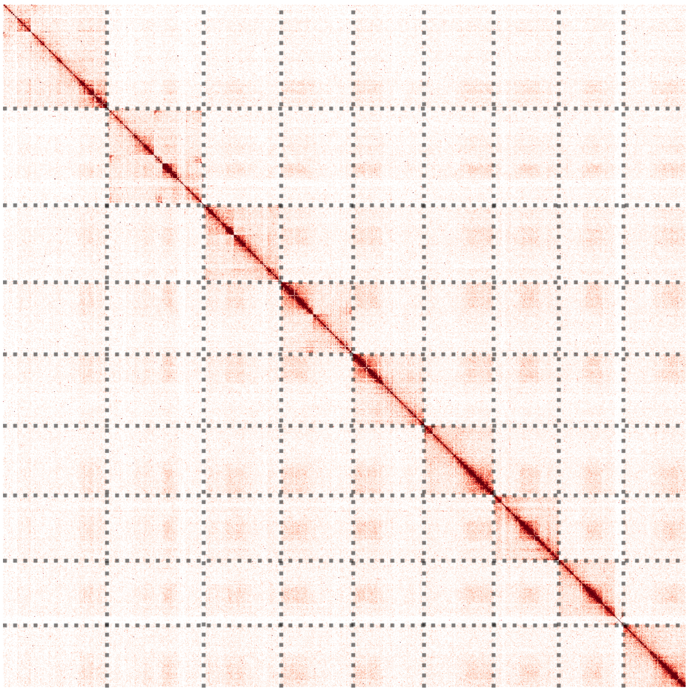

**Fig. S1**

Hi-C contact map of *Platynothrus peltifer* with a binning of 500 representing 9 chromosome-level scaffolds (chromosome 1 to 9 from left to right, top to bottom). The contact map shows strong intrachromosomal interaction frequencies and no structural errors.

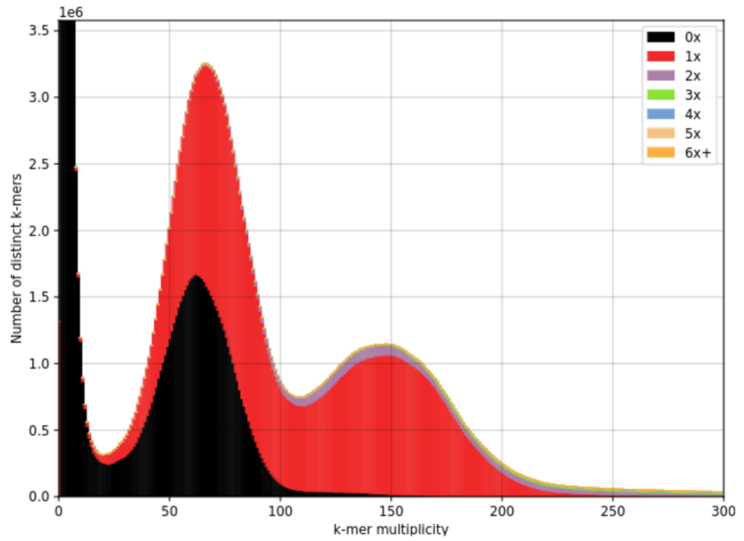


(a) Collapsed assembly Hap0.

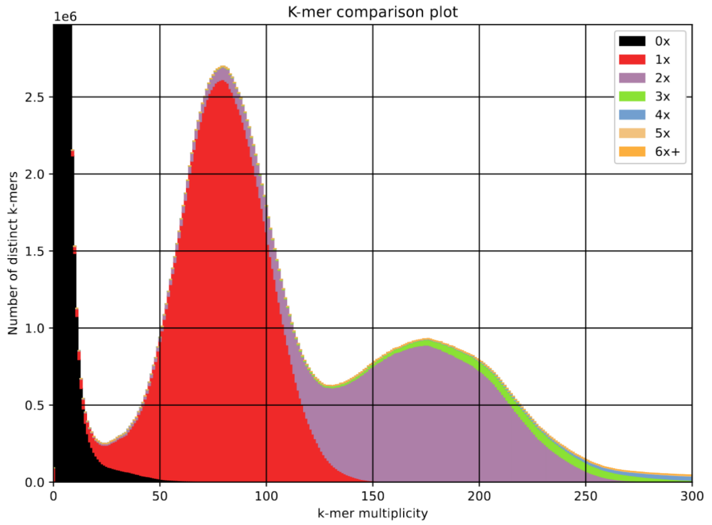


(b) Phased assembly.

**Fig. S2.**

*k*-mer comparison of the assemblies of *Platynothrus peltifer* against PacBio HiFi reads. The plots show two peaks, the first one for heterozygous *k*-mers, and the second one for homozygous *k*-mers. In both assemblies, low-multiplicity *k*-mers are not included (black, 0X). a) Final chromosome-level collapsed assembly. Homozygous *k*-mers are represented exactly once (red). As the assembly is collapsed, only one version of each heterozygous region can be included in the assembly, thus part of heterozygous *k*-mers are represented once (red) and part are absent from the assembly (black). b) Phased assembly including haplotype blocks A and B. Homozygous *k*-mers are represented twice (purple) for both haplotypes and heterozygous *k*-mers are all represented once (red).

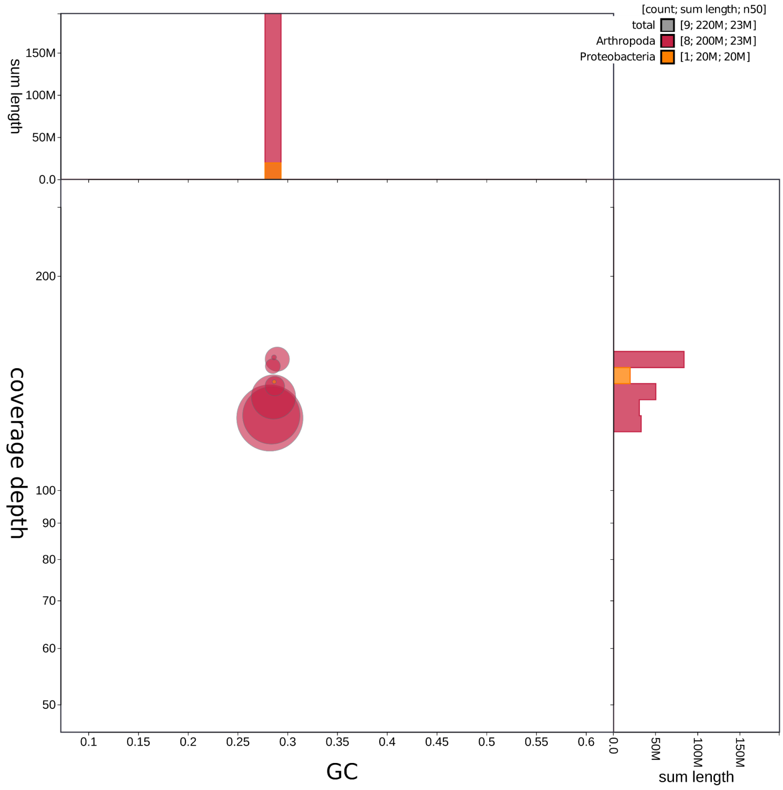


**Fig. S3.**

The final assembly (Hap0 scaffolds) contains no contaminants as seen in the blob plot analysis. The nine chromosome candidates have similar GC contents of 0.28, and their coverage depth (based on mapping of the HiFi reads) ranges from 126X to 153X. Eight of these scaffolds were flagged as Arthropoda, and chr8 was flagged as Proteobacteria due to the integration of *Wolbachia* sequences in the genome.

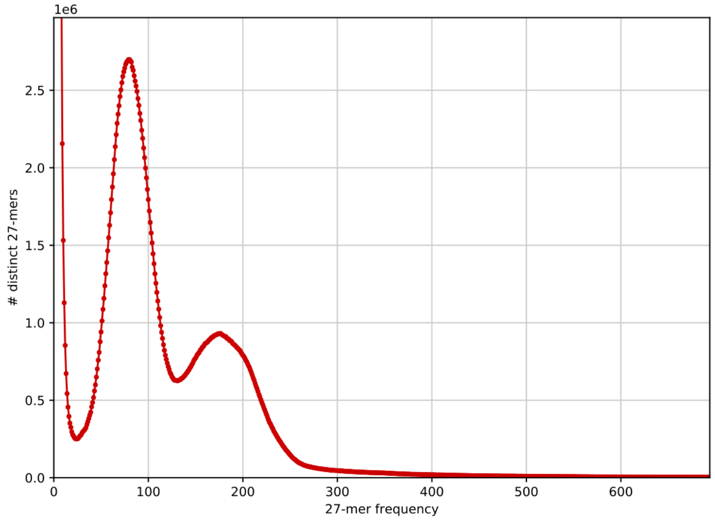


(a) 27-mer histogram.

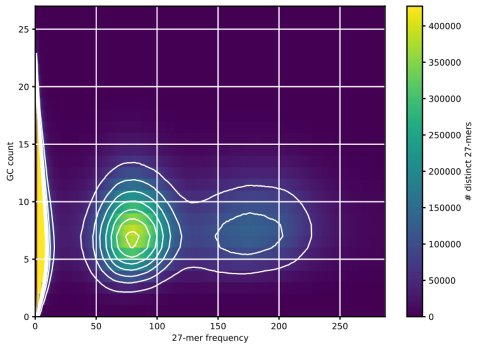


(b) 27-mer GC content analysis.

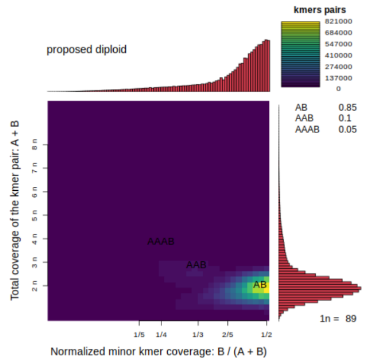


(c) Smudgeplot

**Fig. S4.**

Genome property analyses based on *k*-mers of the PacBio HiFi reads ($k = 27$). a) In the *k*-mer histogram, two peaks are visible at 80X and 160X corresponding to the heterozygous *k*-mers and the homozygous *k*-mers respectively, which suggests a diploid, heterozygous genome. b) The two peaks identified in histogram (a) have similar GC content, and there is no extra peak at a higher GC content, which indicates that there is no significant contamination in the reads. c) The Smudgeplot shows a distinct smudge of *k*-mers with an A/B configuration and the genome is identified as diploid without duplications.

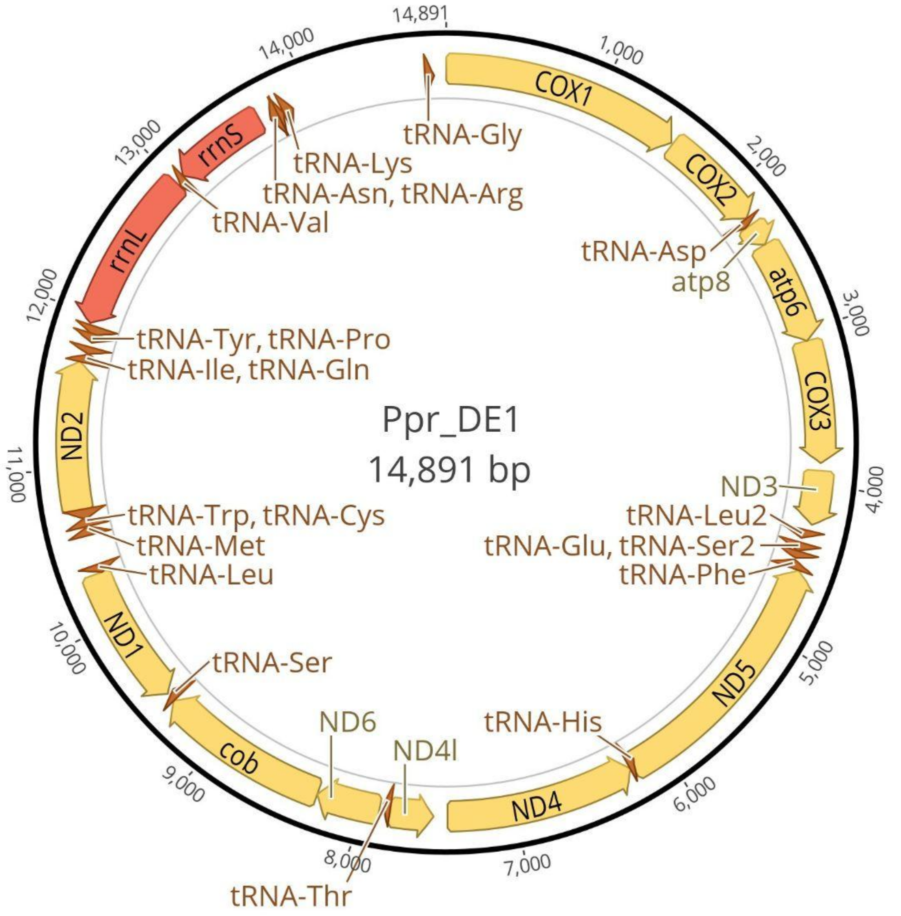


**Fig. S5.**

Annotated mitochondrion of *Platynothrus peltifer*, stemming from the same individual as the reference genome. The size is 14,891 bp and all typical mitochondrial genes could be identified.

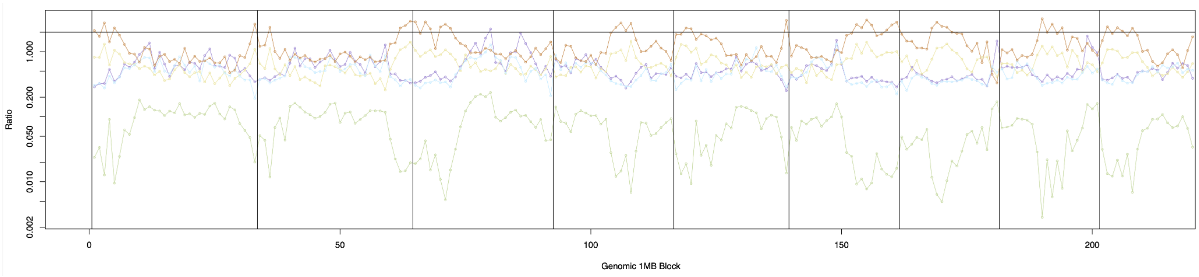


**Fig. S6.**

The ratio of unshared to shared heterozygotic sites from pairwise comparisons, averaged for each population, and separated into 1 Mb genomic blocks. The expected value of this ratio is 2 (black line) in a sexual, random mating population, and trends towards 0 under long-term clonal reproduction. In a sexual population, the expected value of Rx = 2. I.e., for each pair of individuals, the ratio should be approximately 2 Het-Hom, 2 Hom-Het, and 1 Het-Het sites. In a clonal asexual population, because the two haplotypes diverge, any site that is either i) het at the beginning, or ii) fixed in one haplotype but not found in the other, will be shared het sites. Unshared het sites only occur when there are newer mutations (in one haplotype) that are not yet fixed. With time, the number of shared het sites will increase indefinitely, but unshared will stay in an equilibrium (new mutations vs fixation/loss within haplotype). Given enough time, Rx will become smaller and smaller. It cannot reach 0 (because there are always new mutations not fixed yet) but should be much smaller than 2.

The proportion of shared heterozygous sites among individuals of each of the populations is much larger for the Italian population, followed by the German, Russian and Japanese, with the Canadian population sharing only few heterozygous sites (fig. S8). As individual heterozygosity values of the European-Russian populations are comparable, the differences likely reflect population size variation, as in smaller populations the likelihood to sample closely related clones of the same lineage increases. The Canadian and Japanese populations have the lowest individual heterozygosity and also fewer shared heterozygous variants, which might suggest more recent transition events to asexuality, much lower mutation rates and/or substantially larger population sizes compared to European-Russian populations.

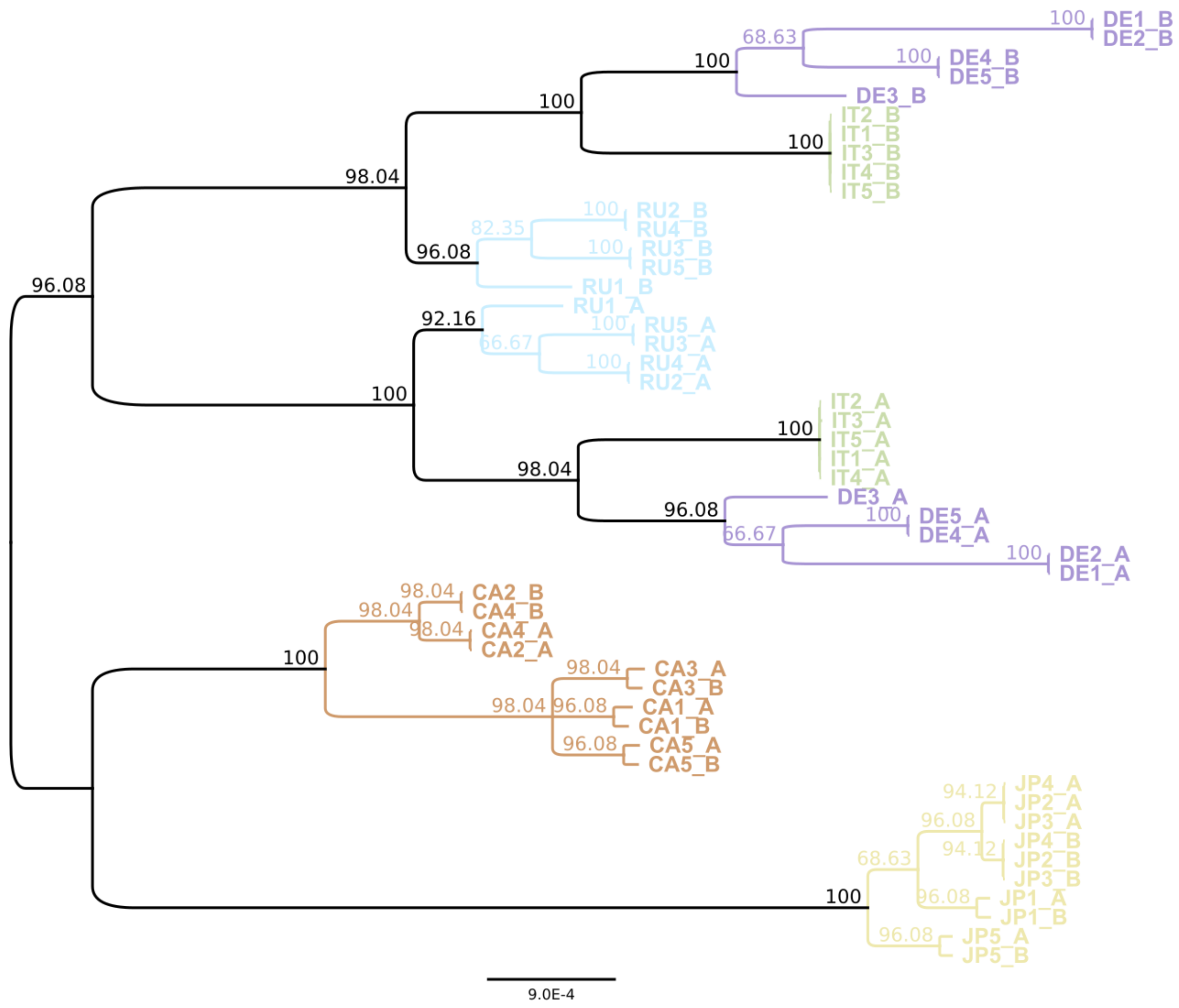


**Fig. S7.**

The haplotype tree illustrates the presence of the Meselson effect in three populations of *Platynothrus peltifer* (Germany - DE, Italy - IT, Russia - RU). In addition, haplotype divergence exceeds divergence among individuals for three individuals from Japan (JA) and two from Canada (CA). The tree is a majority rule consensus tree (i.e. comprises clades that are present in at least 50% of trees) constructed from 51 trees. Each of the 51 trees was reconstructed from a 1 Mb bin derived from the longest alignment block of the in-phase regions per chromosome. For readability the tree was rooted manually at the clade comprising the populations DE, IT, RU. Branch labels represent consensus support values (%; e.g. a value of 96.08 indicates that a clade is present in 96.08% (49) of 51 trees). Branch length indicates divergence (average number of substitutions per site). Colors indicate different populations; _A & _B: haplotypes A and B.

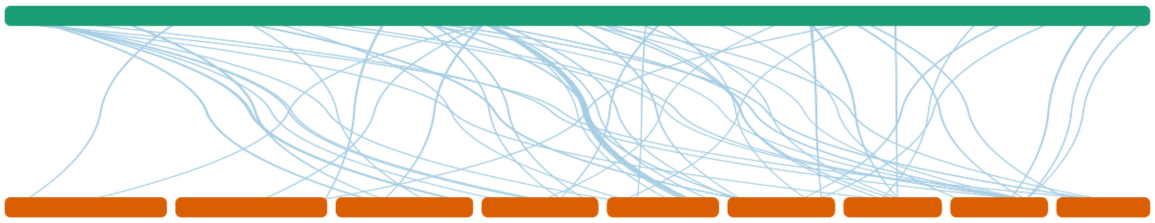

**Fig. S8.**

Integration of *Wolbachia* remnants into the *Platynothrus peltifer* genome. Depending on the stringency of parameters, portions of *Wolbachia* can be detected. Synteny analyses suggest a *Wolbachia* genome (in green) copies throughout the *P. peltifer* genome (in red).

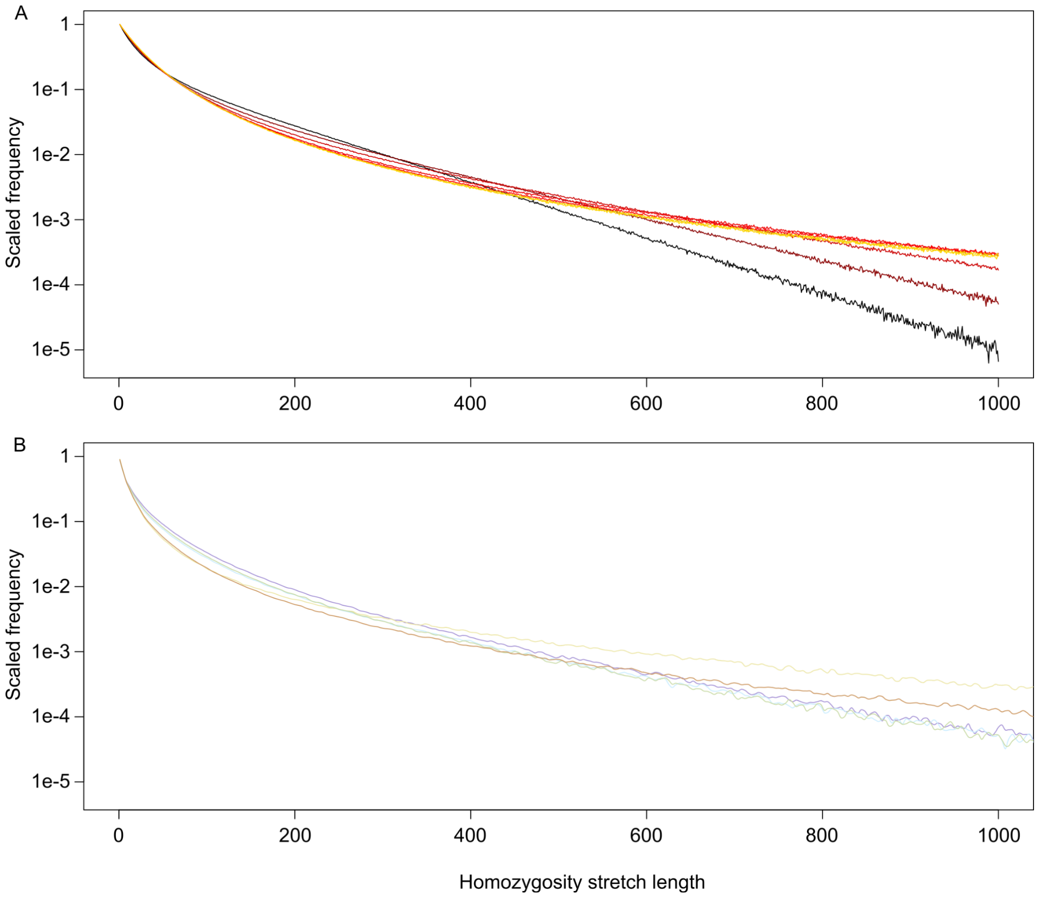


**Fig. S9.**

Gene conversion exists in *Platynothrus peltifer* and the track lengths are at least 500bp. Distribution of homozygosity stretch lengths in (A) simulated and (B) empirical data. The numbers are normalized by dividing with the counts of stretches of size 0 (consecutive sites are heterozygous). The simulation shows the effects of GC track length on the 'homozygosity stretch', i.e. number of consecutive matching nucleotides between haplotypes. If the process is governed by mutation alone, each site should be independent, and lengths of homozygosity stretches geometrically distributed. GC biases the distribution so that longer stretches are more likely to occur (*76*). Line colors in (A) represent different GC track lengths: black = 50bp; dark to light red = 100, 200, 500, 1000bp; dark yellow and yellow = 2000, 5000bp and in (B) empirical mean track length estimates of the different populations. When the mean GC track is short (50bp), the distribution resembles the expected geometric; with the increase of track length, it becomes flatter by reducing the number of stretches < 400bp and increasing those > 400bp. However, once the mean GC track is 500bp or longer, increasing track length does not seem to change the distribution anymore. This asymptotic distribution is highly similar to the one observed from the empirical *P. peltifer* data.

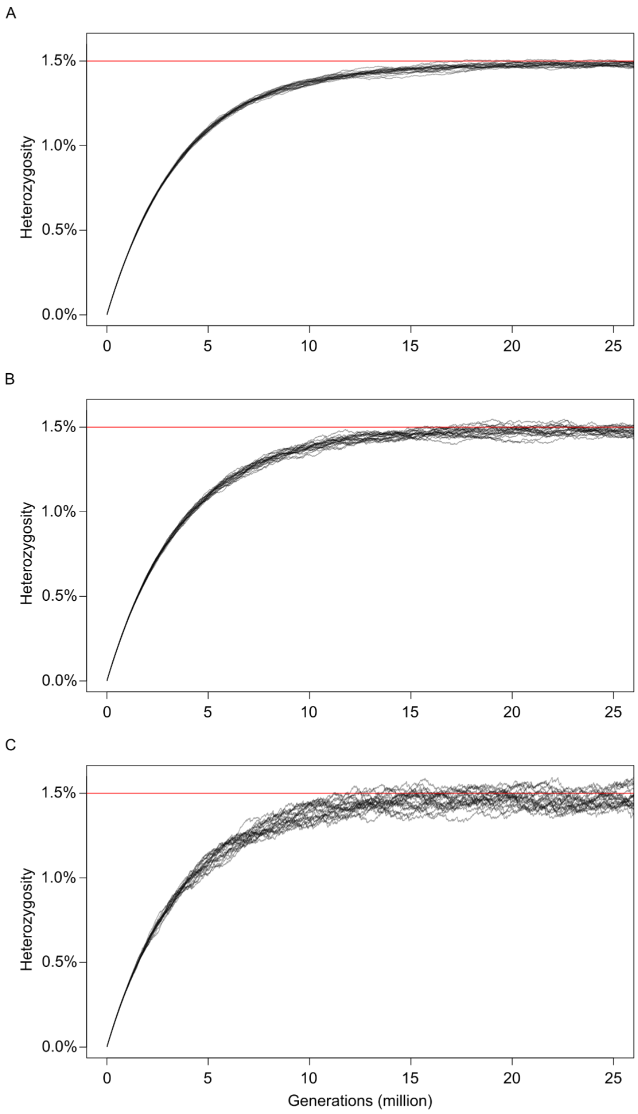


**Fig. S10.**

Simulations of heterozygosity change (i.e., divergence of haplotypes) over time under different gene conversion track length in *Platynothrus peltifer*. The simulation was started from a hypothetical zero-heterozygosity pair of haplotypes, and the mutation rates estimated from *P. peltifer* and a gene conversion rate that gives a theoretical equilibrium heterozygosity of 1.5% (mean genome-wide heterozygosity of five populations) were applied. Average number of nucleotides affected by gene conversion per event were A: 200 B: 1000 C: 5000 bp. While a longer GC track does increase the variation of heterozygosity, it has a small effect compared to the total heterozygosity value.

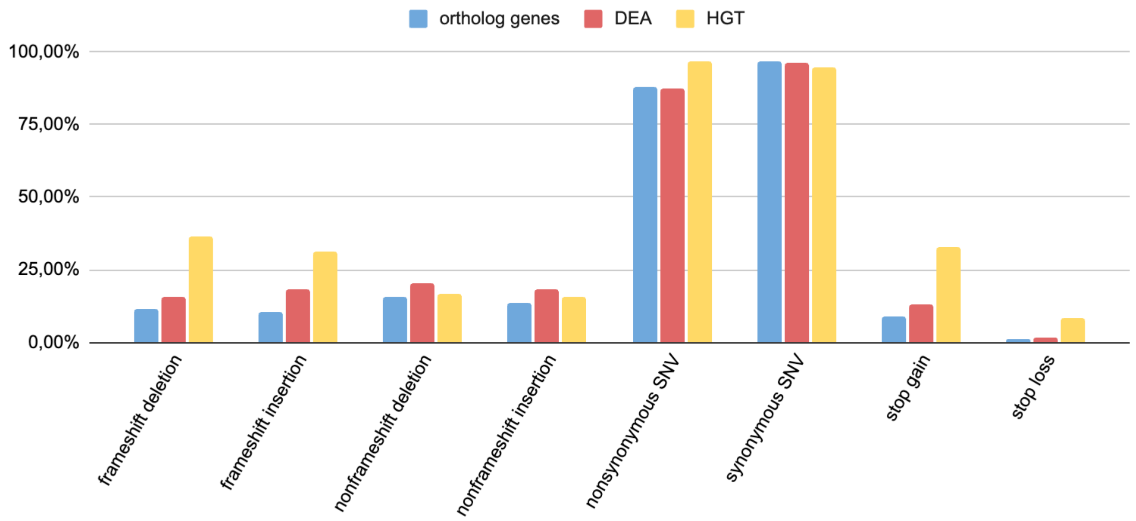


**Fig. S11.**

Categories of different SNP effects and their percentage in orthologous genes (blue), DEAs (red) and HGTs (yellow).

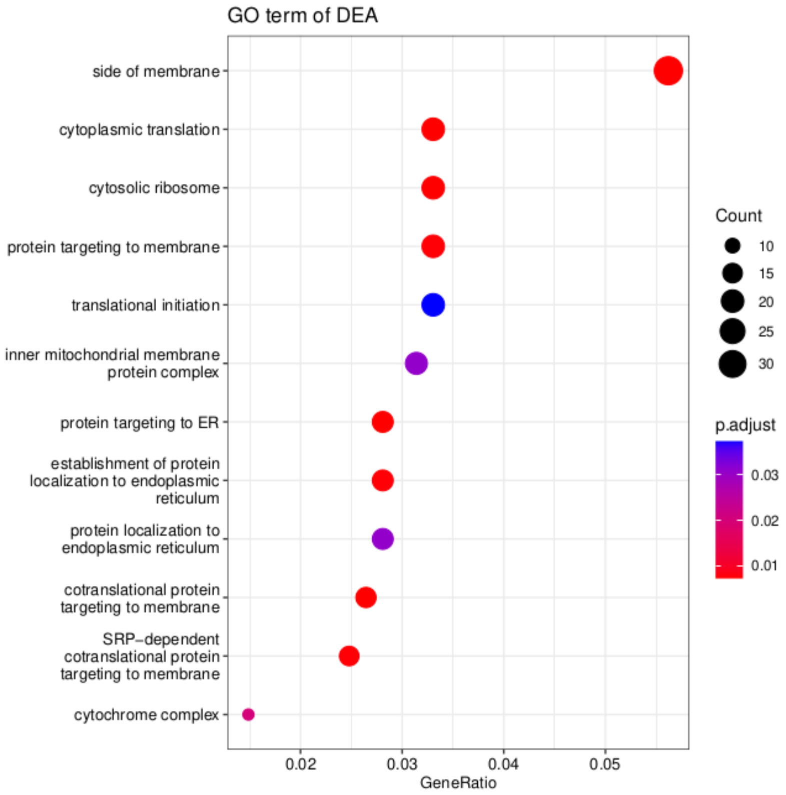


**Fig. S12.**

Enrichment of differentially expressed alleles (DEAs) in GO terms.

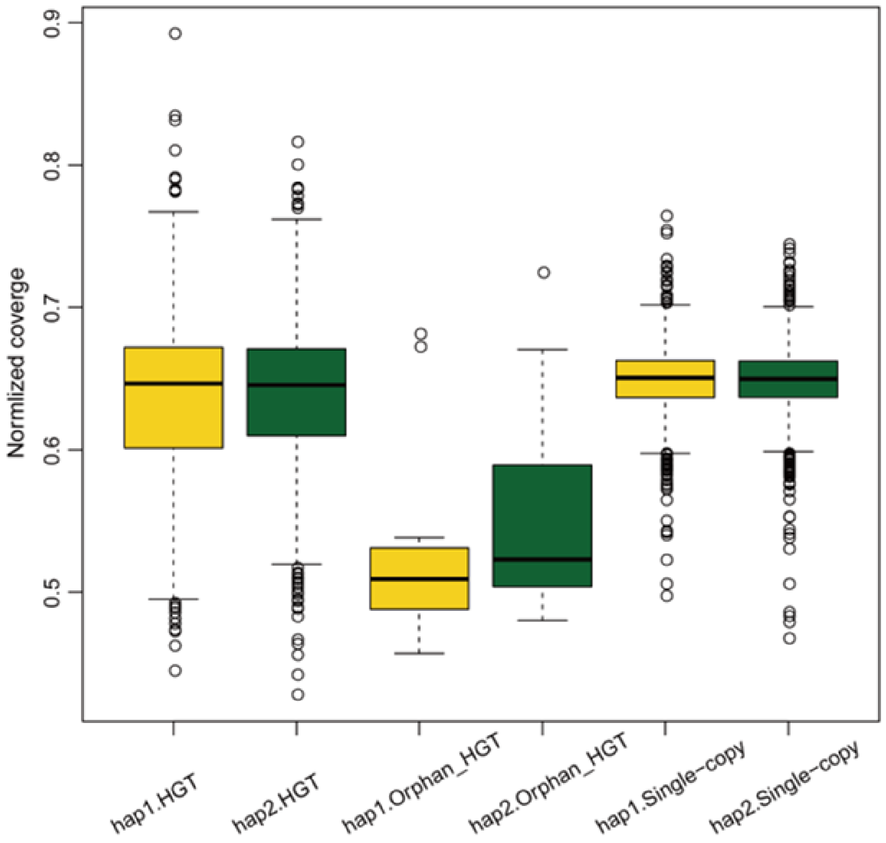


**Fig. S13.**

The orphan HGTs show reduced mapped read coverage compared to HGTs and BUSCO single-copy genes found on both haplotypes. This is consistent with annotations of orphan HGTs being present in only one haplotype and suggests no missing HGT allele in the haplotype assemblies.

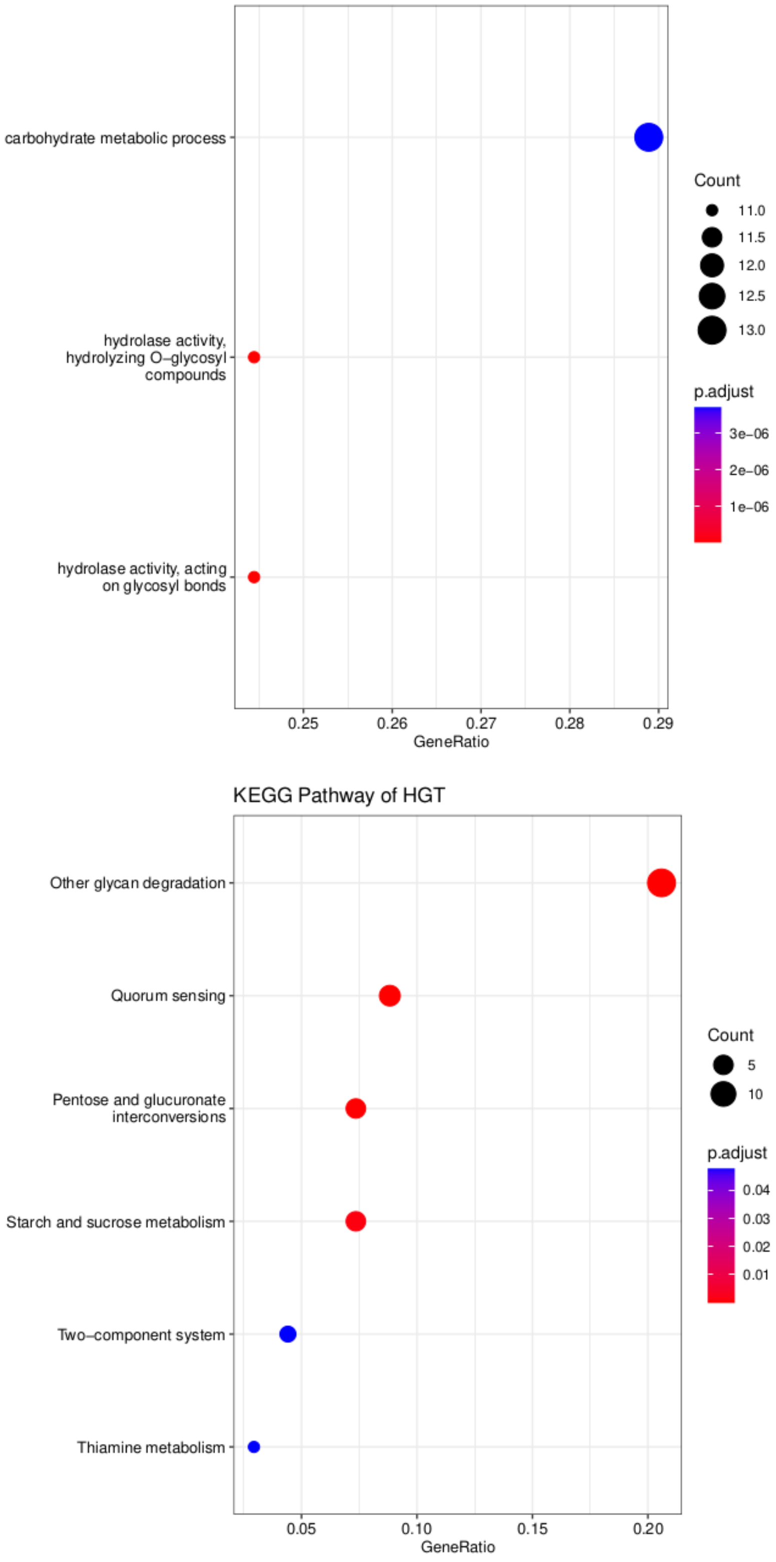


**Fig. S14.**

Enrichment of HGTs in a) GO terms and b) KEGG pathways.

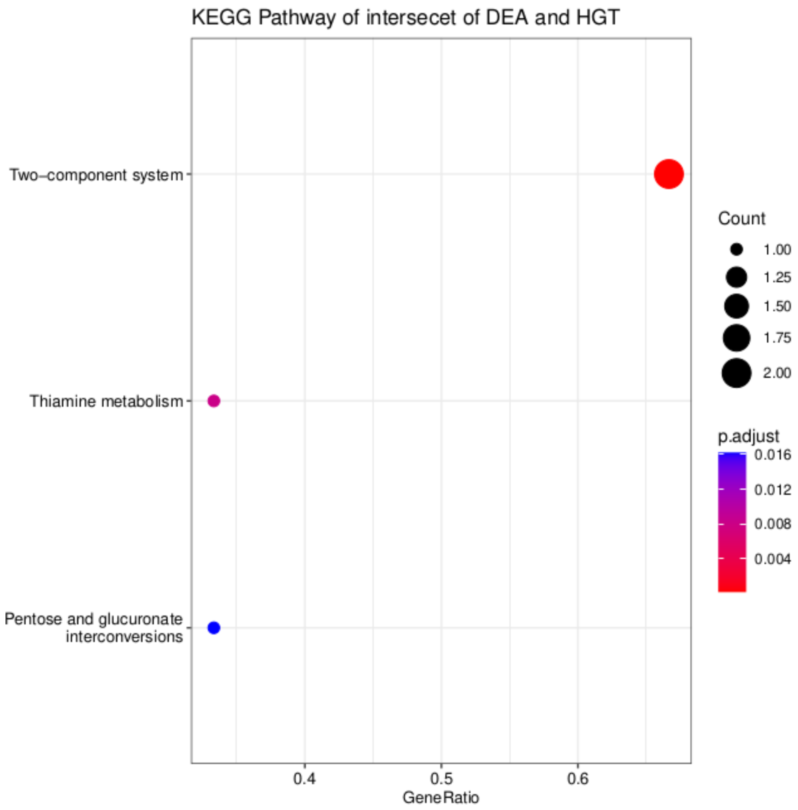


**Fig. S15.**

KEGG pathways enrichment of HGT genes that exhibit differential expression.

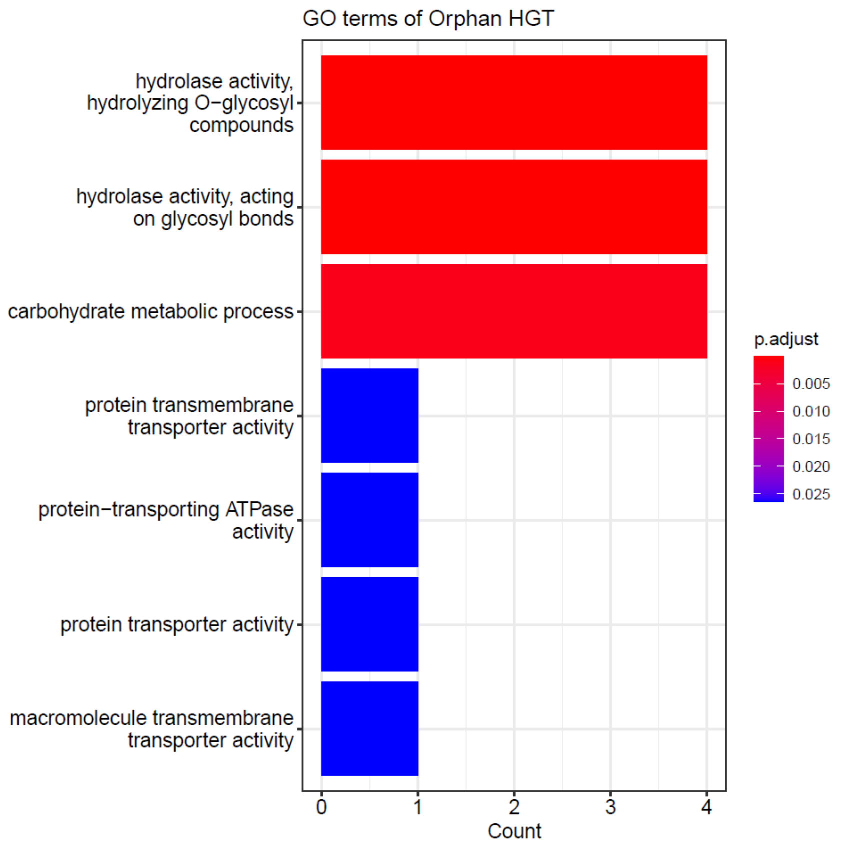


**Fig. S16.**

GO term enrichment for orphan HGTs.

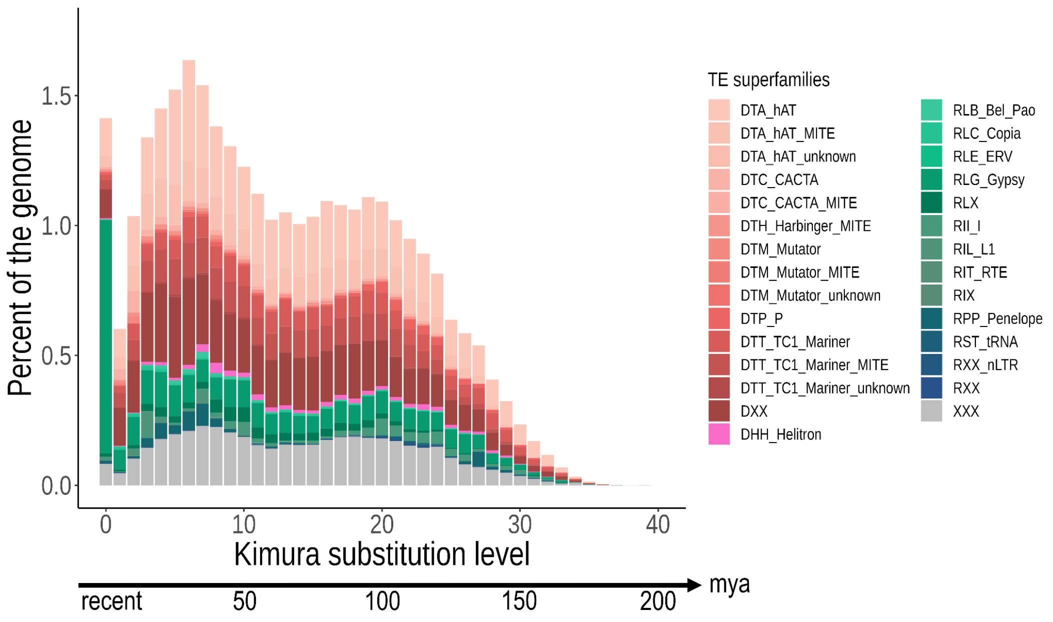


**Fig. S17.**

Transposable element (TE) content and activity in the *Platynothrus peltifer* reference genome. The TE landscape shows historical TE activity and indicates recent activity and a main past expansion event approximately 29-49 mya (6%-10% divergence), coinciding with increase of global temperature dynamics.

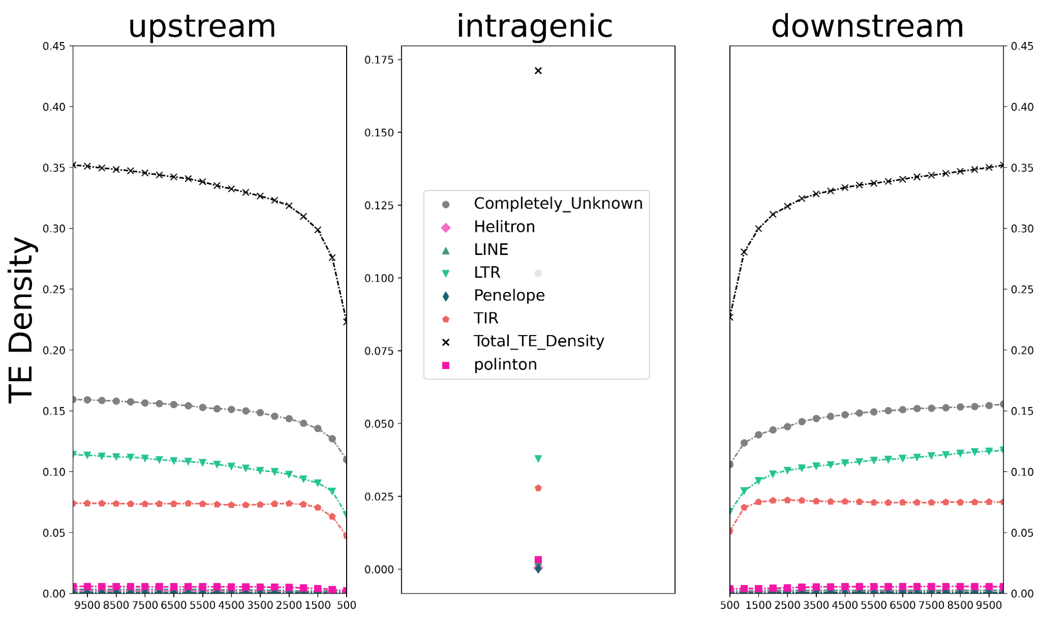


**Fig. S18.**

Transposable elements are selected against in the *Platynothrus peltifer* reference genome. TE density patterns (of chromosome 1) upstream, intragenic and downstream suggest effective selection against TE insertions in the proximity and within genes (note differences in scale).

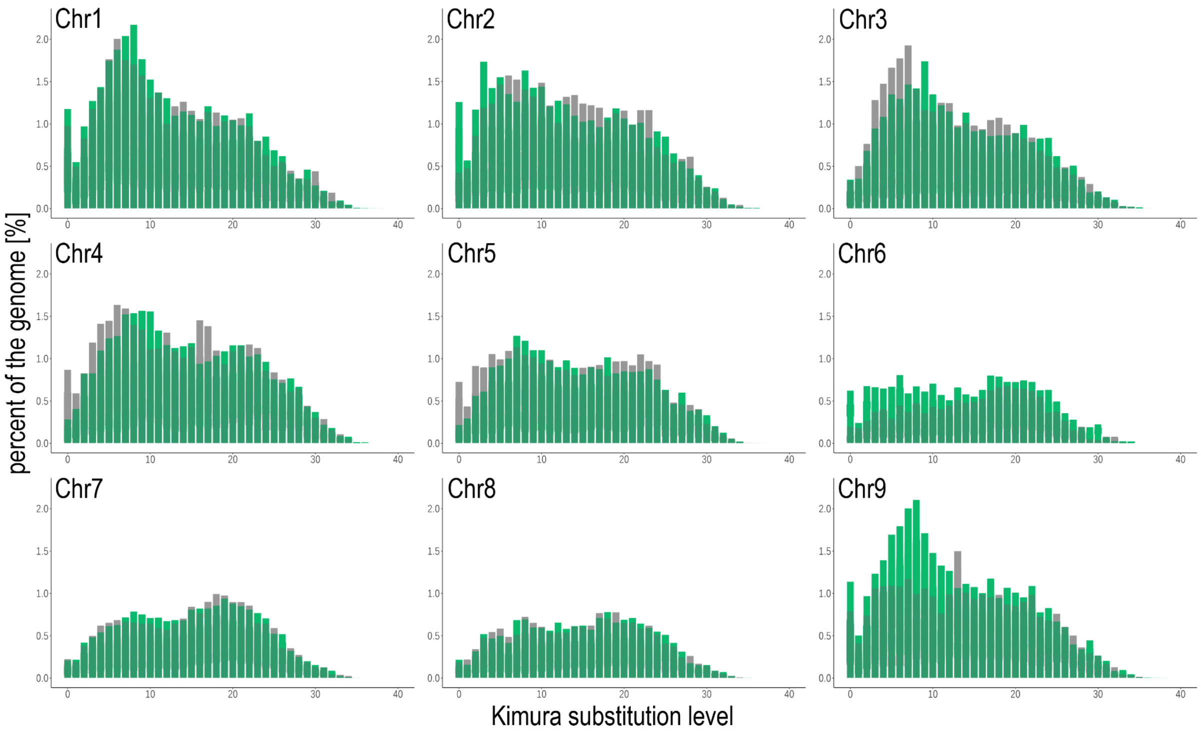


**Fig. S19.**

Transposable element divergence landscapes of the largest haplotypic blocks per chromosome show slight but noticeable haplotype specific historical activity. Green bars represent haplotypic blocks A and gray bars haplotypic blocks B. For example, chr2 and chr5 show haplotype-specific repeat dynamics with a pronounced divergence in activity approximately at 6%, which translates into 29 mya. For chr4, chr6 and chr9 activity differences starting from 12% divergence translate to 59 mya.

**Table S1.**

Assembly and annotation metrics of *Platynothrus peltifer*.

| Metrics | Value |
|---|---|
| Assembly size [Mb] | 219 |
| N50 [Mb] | 23 |
| Number of scaffolds | 9 |
| BUSCO completeness [%] (arthropoda_odb10) | C:96.0 [S:93.9, D:2.1], F:1.0, M:3.0 |
| BUSCO completeness [%] (arachnida_odb10) | C:96.7 [S:93.7, D:3.0], F:1.0, M:2.3 |
| Nr annotated genes | 24.9 |
| Nr annotated genes (both haplotypes) | 10.24 |
| Repeats [%] | 31.9 |
| BUSCO Protein [%] | 98 |

**Table S2.**

Fragment sizes [bp] of haplotypic blocks A and B (in bold the longest scaffold in each chromosome used for Meselson effect and TE analyses)

| Haplotypic blocks A | | Haplotypic Blocks B | |
|---|---|---|---|
| **alt1_chr1_1** | **14957471** | **alt2_chr1_1_1** | **16050112** |
| alt1_chr1_2 | 3366384 | alt2_chr1_1_2 | 30068 |
| alt1_chr1_3 | 11361060 | alt2_chr1_2_1 | 3329658 |
| | | alt2_chr1_3_1 | 11289522 |
| alt1_chr2_1 | 4591776 | alt2_chr2_1_1 | 4510490 |
| alt1_chr2_2 | 5547886 | alt2_chr2_1_2 | 30298 |
| alt1_chr2_3 | 7322949 | alt2_chr2_2_1 | 5700626 |
| alt1_chr2_4 | 5292 | alt2_chr2_3_1 | 9480104 |
| **alt1_chr2_5** | **10752214** | alt2_chr2_3_2 | 163756 |
| alt1_chr2_6 | 7574 | alt2_chr2_3_3 | 41987 |
| alt1_chr2_7 | 46558 | **alt2_chr2_5_1** | **8601668** |
| | | alt2_chr2_5_2 | 47936 |
| alt1_chr3_1 | 676958 | alt2_chr3_10_1 | 437948 |
| alt1_chr3_2 | 3879045 | alt2_chr3_10_2 | 34120 |
| alt1_chr3_3 | 5119837 | alt2_chr3_11_1 | 126106 |
| alt1_chr3_4 | 38148 | alt2_chr3_1_1 | 657132 |
| **alt1_chr3_5** | **12829885** | alt2_chr3_2_1 | 4292554 |
| alt1_chr3_6 | 16941 | alt2_chr3_2_2 | 26733 |
| alt1_chr3_7 | 1421540 | alt2_chr3_2_3 | 23797 |
| alt1_chr3_8 | 22959 | alt2_chr3_2_4 | 15897 |
| alt1_chr3_9 | 25189 | alt2_chr3_3_1 | 4992518 |
| alt1_chr3_10 | 375324 | alt2_chr3_4_1 | 35663 |
| alt1_chr3_11 | 179310 | **alt2_chr3_5_1** | **11108905** |
| alt1_chr3_12 | 16590 | alt2_chr3_5_2 | 115854 |
| alt1_chr3_13 | 17503 | alt2_chr3_6_1 | 16643 |
| | | alt2_chr3_7_1 | 966844 |
| | | alt2_chr3_7_2 | 21497 |
| | | alt2_chr3_8_1 | 38849 |
| | | alt2_chr3_9_1 | 47801 |
| alt1_chr4_1 | 4929078 | alt2_chr4_1_1 | 5114047 |
| alt1_chr4_2 | 3965152 | alt2_chr4_2_1 | 3867448 |
| alt1_chr4_3 | 4381018 | alt2_chr4_2_2 | 813 |

| | | | |
|---|---|---|---|
| **alt1_chr4_4** | **9443682** | alt2_chr4_2_3 | 17851 |
| alt1_chr4_5 | 6884 | alt2_chr4_2_4 | 32058 |
| | | alt2_chr4_3_1 | 6073 |
| | | alt2_chr4_3_2 | 4246053 |
| | | **alt2_chr4_4_1** | **9773924** |
| | | alt2_chr4_4_2 | 42082 |
| **alt1_chr5_1** | **13159702** | **alt2_chr5_1_1** | **12553406** |
| alt1_chr5_2 | 3558262 | alt2_chr5_1_2 | 901490 |
| alt1_chr5_3 | 4240044 | alt2_chr5_2_1 | 2145778 |
| | | alt2_chr5_3_1 | 5744347 |
| alt1_chr6_1 | 6313945 | alt2_chr6_1_1 | 6077717 |
| alt1_chr6_2 | 3497893 | alt2_chr6_2_1 | 2259272 |
| alt1_chr6_3 | 516618 | alt2_chr6_3_1 | 476161 |
| alt1_chr6_4 | 32118 | alt2_chr6_4_1 | 16509 |
| **alt1_chr6_5** | **7722251** | **alt2_chr6_5_1** | **9189643** |
| alt1_chr6_6 | 2805762 | alt2_chr6_5_2 | 11517 |
| | | alt2_chr6_5_3 | 32350 |
| | | alt2_chr6_6_1 | 2871417 |
| **alt1_chr7_1** | **15067032** | **alt2_chr7_1_1** | **15422707** |
| alt1_chr7_2 | 3480545 | alt2_chr7_1_2 | 18765 |
| | | alt2_chr7_1_3 | 15977 |
| | | alt2_chr7_2_1 | 3041516 |
| **alt1_chr8_1** | **10309134** | **alt2_chr8_1_1** | **10145487** |
| alt1_chr8_2 | 7338071 | alt2_chr8_1_2 | 20432 |
| alt1_chr8_3 | 321376 | alt2_chr8_2_1 | 8024092 |
| | | alt2_chr8_3_1 | 268508 |
| **alt1_chr9_1** | **17136559** | **alt2_chr9_1_1** | **16549588** |
| alt1_chr9_2 | 1763873 | alt2_chr9_1_2 | 71763 |
| No_1 | 29782 | alt2_chr9_1_3 | 28075 |
| | | alt2_chr9_1_4 | 27189 |
| | | alt2_chr9_2_1 | 2038438 |

**Table S3.**

Summary statistics for the longest in-phase region that was used for haplotype tree reconstruction per chromosome. Target start and target end mark the start and end coordinates to extract the longest alignment block

| query name | query length | query start | query end | strand direct-ion | target name | target length | target start | target end | # residue matches | alignment block length |
|---|---|---|---|---|---|---|---|---|---|---|
| alt1_chr1_1 | 14957471 | 3454174 | 9999998 | + | chr1 | 32608290 | 3587612 | 10326678 | 5228739 | 6864089 |
| alt1_chr2_5 | 10752214 | 28598 | 5717847 | + | chr2 | 30492103 | 89265 | 5854202 | 4686177 | 5862286 |
| alt1_chr3_3 | 5119837 | 32321 | 5119734 | + | chr3 | 27587832 | 45418 | 5170894 | 4366896 | 5196015 |
| alt1_chr4_1 | 4929078 | 15814 | 4928941 | + | chr4 | 23397169 | 14530458 | 19553083 | 4185504 | 5063857 |
| alt1_chr5_1 | 13159702 | 2582264 | 6054325 | + | chr5 | 22442908 | 4686733 | 8214567 | 3064880 | 3552017 |
| alt1_chr6_5 | 7722251 | 1802320 | 7722233 | + | chr6 | 21557542 | 13485156 | 19401970 | 5149241 | 6050008 |
| alt1_chr7_1 | 15067032 | 7073018 | 12787819 | + | chr7 | 19673716 | 7426596 | 13193208 | 4927722 | 5883881 |
| alt1_chr8_1 | 10309134 | 3280164 | 7993029 | + | chr8 | 19511964 | 8588947 | 13277971 | 4100966 | 4785435 |
| alt1_chr9_1 | 17136559 | 7136559 | 11645221 | + | chr9 | 18765566 | 7084104 | 11663639 | 3571460 | 4657571 |

**Table S4.**

Genomic proportion of different transposable element superfamilies detected in the genome.

| **TE superfamily** | Proportion in genome |
|---|---|
| DHH_Helitron | 0.447 |
| DTA_hAT | 6.451 |
| DTA_hAT_MITE | 2.737 |
| DTA_hAT_unknown | 0.071 |
| DTC_CACTA | 0.388 |
| DTC_CACTA_MITE | 0.075 |
| DTH_Harbinger_MITE | 0.243 |
| DTM_Mutator | 0.074 |
| DTM_Mutator_MITE | 0.097 |
| DTM_Mutator_unknown | 0.004 |
| DTP_P | 0.481 |
| DTT_TC1_Mariner | 2.173 |
| DTT_TC1_Mariner_MITE | 2.618 |
| DTT_TC1_Mariner_unknown | 0.111 |
| DXX | 5.570 |
| RII_I | 0.599 |
| RIL_L1 | 0.214 |
| RIT_RTE | 0.026 |
| RIX | 0.005 |
| RLB_Bel_Pao | 0.152 |
| RLC_Copia | 0.139 |
| RLE_ERV | 0.033 |
| RLG_Gypsy | 3.339 |
| RLX | 0.706 |
| RPP_Penelope | 0.494 |
| RST_tRNA | 0.015 |
| RXX | 0.038 |
| RXX_nLTR | 0.081 |
| XXX | 4.551 |
| **Total TEs in genome** | 31.932 |
